# Roles and interactions of the specialized initiation factors EIF4E2, EIF4E5 and EIF4E6 in *Trypanosoma brucei*: EIF4E2 maintains the abundances of S-phase mRNAs

**DOI:** 10.1101/2022.05.10.491326

**Authors:** Franziska Falk, Rafael Melo Palhares, Albina Waithaka, Christine Clayton

**Affiliations:** Heidelberg University Centre for Molecular Biology (ZMBH), Im Neuenheimer Feld 282, D69120 Heidelberg, Germany

**Keywords:** eukaryotic translation initiation factor 4E, mRNA decay, cell cycle, Pumilio

## Abstract

*Trypanosoma brucei* has six versions of the cap-binding translation initiation factor EIF4E. We investigated the functions of EIF4E2, EIF4E3, EIF4E5 and EIF4E6 in bloodstream forms. We confirmed the protein associations previously found in procyclic forms, and detected specific co-purification of some RNA-binding proteins. Bloodstream forms lacking EIF4E5 grew normally and differentiated to replication-incompetent procyclic forms. Depletion of EIF4E6 inhibited bloodstream-form trypanosome growth and translation. EIF4E2 co-purified only the putative RNA binding protein SLBP2. Bloodstream forms lacking EIF4E2 multiplied slowly, had a low maximal cell density, and expressed the stumpy-form marker PAD1, but showed no evidence for enhanced stumpy-form signalling. EIF4E2 knock-out cells differentiated readily to replication-competent procyclic forms. EIF4E2 was strongly associated with mRNAs that are maximally abundant in S-phase, three of which are bound and stabilized by the Pumilio domain protein PUF9. The same mRNAs had decreased abundances in EIF4E2 knock-out cells. Yeast 2-hybrid results suggested that PUF9 interacts directly with SLBP2, but PUF9 was not detected in EIF4E2 pull-downs. We suggest that the EIF4E2-SLBP2 complex might interact with PUF9, and its bound RNAs, only early during G1/S, stabilizing the mRNAs in preparation for translation later in S-phase or in early G2.

## Introduction

The mechanism of eukaryotic translation initiation has been studied almost exclusively in Opisthokonts, in particular animal cells and *Saccharomyces cerevisiae* (reviewed in (Shirokikh & Preiss, 2018)). In these organisms, the first step is usually binding of the translation initiation factor eIF4E to the mRNA cap. eIF4E is joined by eIF4G, which binds to the helicase eIF4A. Meanwhile, small ribosomal subunits, together with a primed Met-tRNA, eIF2, eIF3, eIF1 and eIF5, together form the 43S complex. The 43S complex is recruited to the cap via an interaction between eIF4G and either eIF3, eIF5, eIF1 or rRNA. The primed small subunit complex scans across the 5’-UTR until it encounters a start codon. Then, a large subunit joins, an exchange of translation factors occurs, and translation can start. The interaction of eIF4E-eIF4G with the cap can be enhanced by interactions between poly(A)-tail bound poly(A) binding protein and eIF4G (Shirokikh & Preiss, 2018). Cap-dependent initiation is, however, possible without eIF4G. For example, in *Arabidopsis*, the CERES protein, which has a canonical eIF4E binding motif but no other eIF4G features, is active in translation initiation through interactions with eIF4A, eIF3 and PABP (Toribio *et al*., 2019). Initiation using a specialized mammalian eIF4G that lacks eIF4E binding has also been demonstrated (de la Parra *et al*., 2018). The efficiency of cap-dependent translation initiation in Opisthokonts can be regulated in numerous ways, for example via the action of proteins that bind the 3’-untranslated region (3’-UTR) and enhance or prevent recruitment of eIF4E/G (Shirokikh & Preiss, 2018). Moreover, most eukaryotes examined so far have more than one paralogue eIF4E and eIF4G. Among the EIF4Es, there is often at least one that has suppressive functions and lacks an eIF4G partner (Hernandez *et al*., 2012).

Trypanosomes and Leishmanias are parasites of animals, and cause a range of human and veterinary diseases. They belong to the class Kinetoplastea, which branched from both Opisthokonts and plants very early in eukaryotic evolution (Burki *et al*., 2020). Leishmanias multiply intracellularly as amastigotes in mammalian hosts, and as promastigotes in sand flies. *Trypanosoma brucei* is an extracellular parasite of mammals, where it multiplies as the bloodstream form, and in Tsetse flies, for which the most commonly studied stage is called the procyclic form. Bloodstream-form *T. brucei* respond to high density by differentiating into a stationary-phase form called the stumpy form, which is pre-adapted for differentiation into the procyclic form (Rico *et al*., 2013). Stumpy forms express a cell surface marker called PAD1 (Dean *et al*., 2009). Bloodstream-form *T. brucei* have a Variant Surface Glycoprotein (VSG) coat which can be switched, enabling the parasites to avoid humoral immunity (Horn, 2014).

Kinetoplastids have six cap-binding EIF4Es (EIF4E1-6) (Dhalia *et al*., 2005, Freire *et al*., 2014a, Freire *et al*., 2014b, Moura *et al*., 2015) and five EIF4Gs (EIF4G1-5) (Dhalia *et al*., 2005, Freire *et al*., 2014a, Freire *et al*., 2014b, Moura *et al*., 2015). Kinetoplastid EIF4E1 has no EIF4G partner, instead binding to two other proteins, 4EIP and 4EIP2. Results so far indicate that *T. brucei* EIF4E1 has primarily a suppressive function (Terrao *et al*., 2018, Falk *et al*., 2021), but additional translation-promoting functions have been suggested for the *Leishmania* homologue (Yoffe *et al*., 2004, Zinoviev *et al*., 2012, Tupperwar *et al*., 2019, Tupperwar *et al*., 2020). One way to investigate the actions of RNA-associated proteins independent of their RNA-binding specificities, is known as “tethering” (Coller & Wickens, 2007). The protein to be investigated is expressed as a fusion with a highly specific RNA-binding domain, such as the RNA-binding peptide of the bacteriophage lambdaN protein. In the same cells, there is a reporter mRNA that includes, in the 3’-UTR, the cognate RNA binding sequence; for lambdaN, this is known as boxB. Both EIF4E1 and 4EIP are strong inhibitors in the tethering assay (Lueong *et al*., 2016, Erben *et al*., 2014).

EIF4E2 (Tb927.10.16070) has no EIF4G partner, instead binding to SLBP2 (Tb927.3.870) (Freire *et al*., 2018), a protein with a domain that, in mammalian cells, binds to the stem-loops at the 3’-termini of histone mRNAs. Kinetoplastid histone mRNAs are - as in most other organisms - polyadenylated and the function of SLBP2 is unknown. In procyclic forms, SLBP2 depletion caused a minor growth defect while depletion of EIF4E2 had no effect on growth or protein synthesis (Freire *et al*., 2011, Freire *et al*., 2018). Tethering of EIF4E2 or SLBP2 fragments to a reporter did not affect expression (Erben *et al*., 2014). Quantitative mass spectrometry results (Supplementary Table S1) suggest that in *T. brucei*, EIF4E2 is 50 times less abundant than EIF4E3 in bloodstream forms, and even less abundant in procyclic forms (Tinti & Ferguson, 2022). *Leishmania* EIF4E2 comigrated with polysomes in sucrose gradients (Yoffe *et al*., 2006), and was much more abundant in promastigotes than in amastigotes.

Until now it has been accepted that EIF4E3 (Tb927.11.11770) and EIF4E4 (Tb927.6.1870) are the major translation initiating EIF4Es in kinetoplastids, because they are roughly equimolar with mRNA (Dhalia *et al*., 2005, Freire *et al*., 2011). *Leishmania* EIF4E3 migrates at 80S (Yoffe *et al*., 2006) and interacts preferentially with EIF4G4 (Moura *et al*., 2015), but *T. brucei* EIF4E3 can interact with both EIF4G3 and EIF4G4 (Freire *et al*., 2011). Deletion of one copy of EIF4E3 in *Leishmania* promastigotes impaired protein synthesis (Shrivastava *et al*., 2019). RNAi targeting EIF4E3 in procyclic and bloodstream form *T. brucei* inhibited growth (Freire *et al*., 2011) and protein synthesis. RNAi targeting *T. brucei* EIF4G3 or EIF4G4 in procyclic forms also impaired growth, with protein synthesis inhibition for EIF4G3 but not EIF4G4 (Moura *et al*., 2015). When EIF4E3, EIF4E4, EIF4G3 or EIF4G4 were tethered to a reporter, they all increased expression (Lueong *et al*., 2016, Erben *et al*., 2014). All of these observations indicate that the EIF4E3/EIF4G4 and EIF4E4/EIF4G3 complexes are active in translation initiation. *Leishmania* EIF4E4 interacts with EIF4G3, which can also bind to 4A1 (Dhalia *et al*., 2005, Yoffe *et al*., 2009), and also directly with PABP1 (Dos Santos Rodrigues *et al*., 2018). Since tagged EIF4E4 was not detected in axenic *Leishmania* amastigotes, it was suggested that another EIF4E might substitute (Zinoviev *et al*., 2011). A more recent publication has shown copurification of *Leishmania* EIF4E4 and EIF4G3 with both PABP1 and an RNA-binding protein, RBP23; and also that both RBP23 and PABP1 were preferentially associated with mRNAs encoding ribosomal proteins (Assis *et al*., 2021). *T. brucei* EIF4E4 also associates preferentially with PABP1, although some association with PABP2 was also detected (Zoltner *et al*., 2018). RNAi targeting EIF4E4 had no effect in *T. brucei* procyclic forms but was lethal in bloodstream forms (Freire *et al*., 2011).

EIF4E5 (Tb927.10.5020) and EIF4E6 (Tb927.7.1670) were described nearly a decade after the other EIF4Es, because they show lower sequence identity and similarity to Opisthokont eIF4Es. The *T. brucei* proteins are now known to be just as abundant as EIF4E3 and EIF4E4 (Dejung *et al*., 2016, Tinti & Ferguson, 2022)(Supplementary Table S1). EIF4E5 is associated with EIF4G1 and EIF4G2 in both *T. brucei* and *Leishmania* (Freire *et al*., 2014b, Shrivastava *et al*., 2021). In *T. brucei*, it was also associated with a dynein light chain (LC8), a putative capping enzyme, the putative RNA-binding protein RBP43, the two 14-3-3 protein paralogues, EF1, and two other proteins of unknown function (Tb927.8.4560, Tb927.11.6010) (Freire *et al*., 2014b). Results suggested that RBP43 and the putative capping enzyme were specific to EIF4G1-containing complexes. EIF4G1 and the interacting proteins were preferentially associated with PABP2, rather than PABP1 (Zoltner *et al*., 2018). RNAi had a very minor effect on procyclic form growth but impaired motility (Freire *et al*., 2014b). Tb927.11.6010 and EIF4G1 both increased expression when tethered, but EIF4G2 did not (Lueong *et al*., 2016); full-length EIF4E5 was not assayed. Interestingly, in *Leishmania*, EIF4E5 binds the cap in promastigotes, but not amastigotes, suggesting that EIF4E5 is required only in the invertebrate host (Shrivastava *et al*., 2021).

In procyclic forms, EIF4E6 pairs with EIF4G5, which in turn interacts with G5-IP (Freire *et al*., 2014a). RNAi in procyclic forms did not affect growth, but did weaken the attachment of the flagellum to the cell body (Freire *et al*., 2014a). Tethered EIF4G5 (Erben *et al*., 2014) or EIF4E6 (Nascimento *et al*., 2020) increase reporter expression, indirectly supporting a role in translation initiation. The EIF4E6/EIF4G5 complex is preferentially associated with the MKT1 complex, which consists of MKT1, PBP1, XAC1 and LSM12 (Nascimento *et al*., 2020), interacts with PABPs, and stimulates translation and RNA stability (Singh *et al*., 2014, Nascimento *et al*., 2020). The MKT1 complex is recruited to mRNAs by RNA-binding proteins, one of which, CFB2, is required to stabilise *VSG* mRNA (Melo do Nascimento *et al*., 2021). EIF4G5 was also weakly co-purified with *VSG* mRNA, suggesting a role in its translation. Recent evidence suggests that *Leishmania* EIF4E6 may have a different role and might even inhibit translation (Tupperwar *et al*., 2021). Although *Leishmania* EIF4E6 also associates with EIF4G5 and G5-IP, the MKT1 and PBP1 homologues did not co-purify (Tupperwar *et al*., 2021). Over-expression of *Leishmania* EIF4E6-SBP inhibited protein synthesis by an unknown mechanism; a hemizygous mutant had morphological defects but normal protein synthesis rates. Unlike *T. brucei* EIF4E6, *Leishmania* EIF4E6 had no detectable binding to the cap analogue m^7^GTP.

Altogether the results so far suggest a repressive role for *T. brucei* EIF4E1, and roles in translation for *T. brucei* EIF4E3-6. We here describe experiments designed to investigate further the functions of EIF4Es 2,5 and 6, comparing with EIF4E3 and focusing particularly on bloodstream-form trypanosomes.

## Results

### EIF4E2, EIF4E3, EIF4E5 and EIF4E6 protein associations in bloodstream forms

The associations of the EIF4Es with other proteins had been tested in procyclic forms, but not in bloodstream forms, and - with the exception of EIF4E1 - had not been analyzed by quantitative mass spectrometry. To re-analyze the associations, we integrated a sequence encoding a Tobacco-Etch-Virus-protease (TEV)-cleavable tag (the PTP tag) at the 3’-ends of the open reading frames, so that each EIF4E would be expressed with the tag at the C-terminus. To be certain that the protein was functional, we then deleted the other allele. This was successful for all proteins except EIF4E4. We therefore focused on EIF4E2, EIF4E5 and EIF4E6, comparing them with EIF4E3 because it was thought to be a general translation initiation factor. Inducibly expressed CFP-PTP served as a general control. We used just a single step of purification, with no RNase treatment, in order to be able to detect proteins with non-stoichiometric association, or which copurified via bound mRNA. The tag contains an IgG domain at the C-terminus, so proteins were first allowed to bind to IgG-beads, then eluted by incubation of the beads with TEV protease. The His-tagged protease was removed using nickel beads before denaturing gel electrophoresis and mass spectrometry. The resulting data were analyzed using MaxQuant, then statistically using Perseus algorithm. The Perseus algorithm substitutes missing values by simulating them, which can lead to artefacts. We therefore eliminated “significantly enriched” proteins that relied primarily on simulated values. Altogether, we detected 1129 different proteins (Supplementary table S2, sheet 2). We initially used 3 replicates for all of the proteins except EIF4E2. Of these, 40 were significantly (p<0.01) at least 4-fold enriched with one or more of EIF4E3, EIF4E5 or EIF4E6. These significantly enriched proteins are listed in Supplementary table S2, sheet 1, and graphs of various comparisons are in Figure 1. In the figure, cut-off curves that apply stricter criteria than the simple “p<0.01, 4x” are also included. Subsequently, three additional replicates for EIF4E2 were analyzed (Supplementary table S2, sheet 3). Supplementary Table S2, sheet 1 includes all proteins that copurified with any of the EIF4Es.

**Figure 1.**
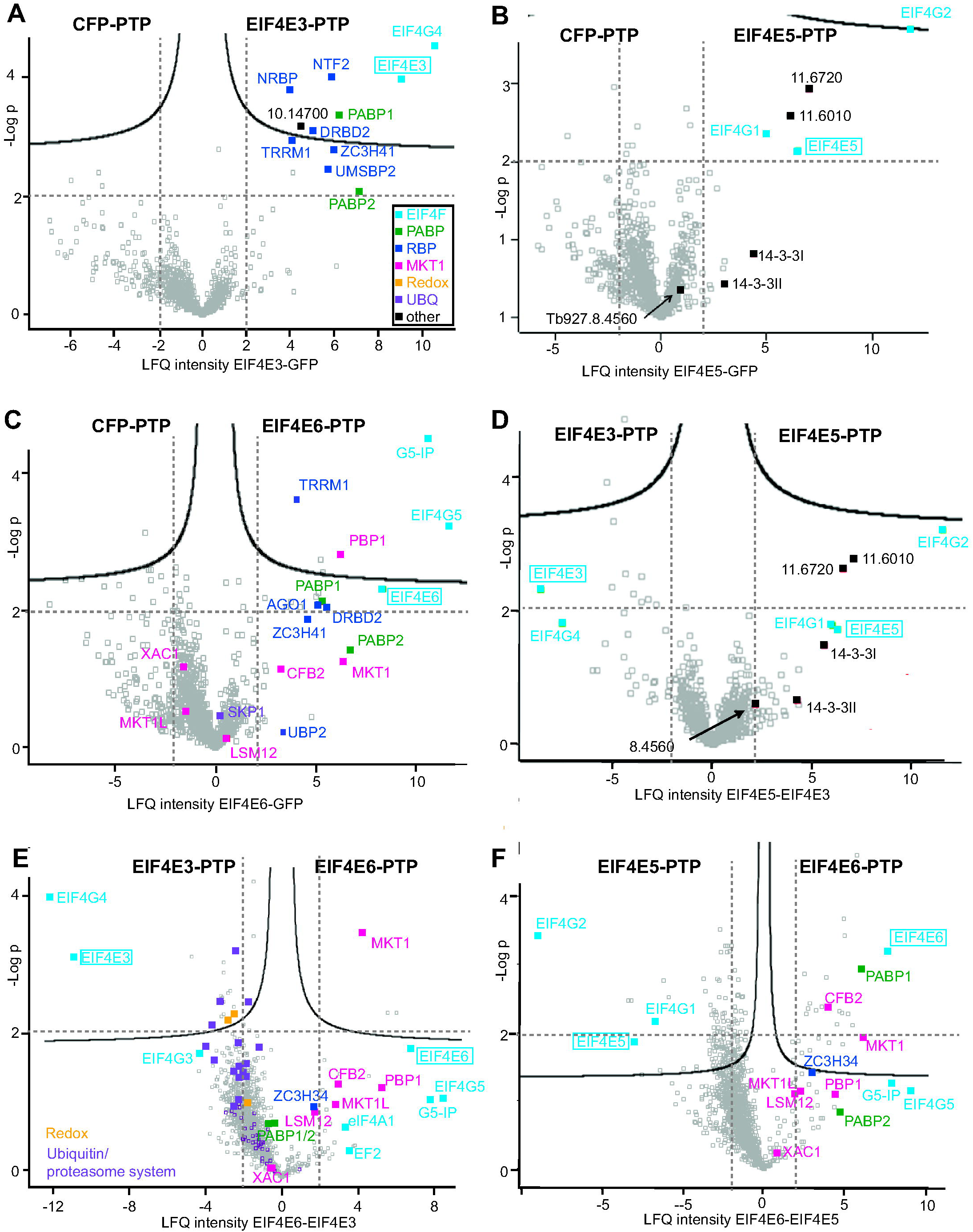
Proteins associated with EIF4E3, 5 and 6. Each protein was expressed from the endogenous locus with a PTP-tag (Schimanski *et al*., 2005) at the C-terminus. The other allele was deleted. Proteins were purified in triplicate on an IgG column and eluted with TEV protease, which was then removed using nickel beads. Inducibly-expressed CFP-PTP was the control. Data were analyzed using Perseus (Tyanova *et al*., 2016), with substitution of absent values. The negative log10 P-value is on the y-axis and the log2 relative enrichment is on the x-axis, with dotted lines indicating 4-fold enrichment and a P-value of 0.01. The continuous lines indicate the significance boundaries estimated by Perseus. Values for selected proteins are highlighted and the label for the PTP-tagged protein is enclosed in a box; gene ID numbers have been shortened by removing the initial “Tb927”. Details are in Supplementary Table S1. **A**. Comparison of EIF4E3-PTP (right) with CFP-PTP (left). The color code for the labels is on the bottom right. RBP: RNA-binding protein; UBQ: ubiquitin-proteasome system. **B**. Comparison of EIF4E5-PTP (right) with CFP-PTP (left). **C**. Comparison of EIF4E6-PTP (right) with CFP-PTP (left). **D**. Comparison of EIF4E5-PTP (right) with EIF4E3-PTP (left). **E**. Comparison of EIF4E6-PTP (right) with EIF4E3-PTP (left). **F**. Comparison of EIF4E6-PTP (right) with EIF4E53-PTP (left).

EIF4E2 copurified only its known partner, SLBP2 (Freire *et al*., 2018) and PABP2 (Supplementary Table S2, sheet 3). Remaining enriched proteins are common contaminants. SLBP2 was previously shown to copurify with PABP2 and PABP1 (Freire *et al*., 2018).

As previously reported, EIF4E3 was strongly associated with EIF4G4 (Supplementary Table S2, Figure 1A). Although EIF4G3 was also detected, it was present in only two of the three replicates and at 100-fold lower intensity. We also detected the poly(A) binding proteins PABP1 and PABP2 in roughly similar amounts. Other proteins that were significantly and specifically enriched were the DNA-binding protein EMSBP2; a nucleolar RNA-binding protein (Tb927.9.5320, NRBP in Figure 1A), a putative guanosine monophosphate reductase (Tb927.5.2080) and the RNA-binding protein ZC3H41 (which was also associated with *VSG* mRNA) (Melo do Nascimento *et al*., 2021). There were also some proteins shared between EIF4E3 and EIF4E6 (Supplementary Table S2, Figure 1A, C): a protein with an NTF2 domain, the DNA-binding protein UMSBP2, DRBD2, TRRM1, MKT1 and Tb927.10.14700. Shared with EIF4E5 (but more abundant with EIF4E3) were an IMP dehydrogenase, the RNA-binding proteins DRBD3/PTB1 and RBP42 and a subunit of the nascent polypeptide associated complex, involved in folding proteins that emerge from the ribosome (Supplementary Table S2).

The EIF4E5 purification (Supplementary Table S2, Figure 1B) contained most of the proteins that had been previously reported to be associated with *T. brucei* EIF4E5 in procyclic forms (Freire *et al*., 2014b), although RBP43 and Tb927.8.4560 were detected in only 2 of the 3 preparations. The homologues of Tb927.11.13060, PUF1, DHH1 and another RNA helicase (Tb927.5.4270) copurified with *Leishmania* EIF4E5 (Shrivastava *et al*., 2021) but were not detected in our purification. We also found EIF4E3 in two of the three EIF4E5 preparations, but not with EIF4E6 or GFP. As previously reported (Zoltner *et al*., 2018) EIF4E5 was associated with PABP2 but not PABP1.

EIF4E6, as expected, co-purified EIF4G5 and the EIF4G5 interacting protein G5-IP (Supplementary Table S2, Figure 1C). MKT1 was clearly more strongly enriched with EIF4E6 than with EIF4E3 (Figure 1E) and CFB2 was found only with EIF4E6.

In summary, our results mainly confirmed that EIF4E associations previously shown in procyclic *T. brucei* are also found in bloodstream forms, and confirmed association of EIF4E6 with the MKT complex and CFB2. The use of quantitative mass spectrometry did however result in specific detection, especially with EIF4E3 and EIF4E6, of a few additional proteins that are known to interact with mRNA.

### EIF4E5 is not essential in cultured bloodstream-form trypanosomes

Depletion of EIF4E5 in procyclic forms slightly inhibited growth and impaired motility, but the extent of depletion after RNAi was not known (Freire *et al*., 2014b). We found that bloodstream-form *T. brucei* completely lacking EIF4E5 (Figure 2A) multiplied at a rate that was within the range of clonal variation (Figure 2B). To test the ability of the cells to differentiate into procyclic forms, we added 6 mM cis-aconitate and dropped the temperature to 27 °C. 24 h later, cells were transferred to procyclic-form medium (MEM-Pros) at 27 °C. The cells expressed the procyclic-form marker EP procyclin with normal kinetics after cis-aconitate addition (Figure 2C) but were unable to multiply as procyclic forms. One clone died immediately and the other lingered for a few days but also eventually died (Figure 2D). These results suggest that EIF4E5 function is more important in procyclic forms than in bloodstream forms.

**Figure 2.**
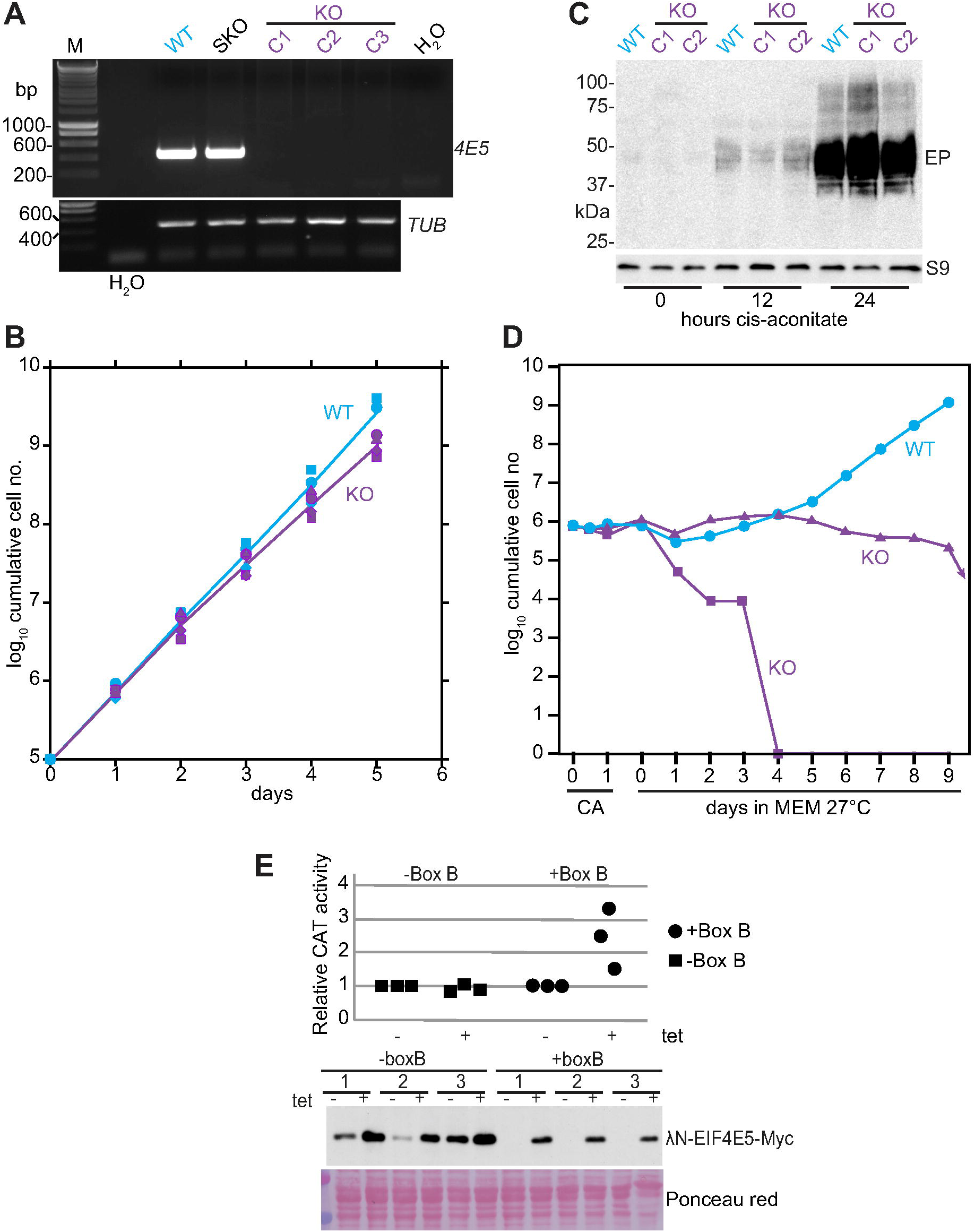
EIF4E5 is not essential in bloodstream forms. Both copies of the EIF4E5 open reading frame were replaced by selectable markers **A**. The *EIF4E5* gene was detected from genomic DNA using internal primers. As a control the beta-tubulin gene was amplified. WT-wild-type; SKO: single knock-out (one gene replaced); KO: double knock-out. **B**. Bloodstream-form *T. brucei* lacking EIF4E5 grow similarly to wild-type. **C**. Bloodstream-form *T. brucei* lacking EIF4E5 express EP procyclin after incubation with *cis*-aconitate at 27 °C. **D**. *T. brucei* lacking EIF4E5 are unable to grow in procyclic-form medium (MEM-Pros) at 27 °C. **E**. EIF4E5 with an N-terminal lambdaN peptide, and a C-terminal myc tag, can activate expression from a reporter mRNA bearing 5 copies of the boxB sequence. The bloodstream-form trypanosomes used contained the chloramphenicol acetyltransferase (*CAT*) reporter mRNA either without (squares) or with (circles) 5 boxB sequences between the open reading frome and the 3’-UTR. lambdaN-EIF4E5-myc was expressed from a tetracycline-inducible promoter, in three different cloned lines per reporter. For each clone the level of CAT enzyme expression in the absence of tetracycline was set to 1. Expression of the myc-tagged protein is shown in the Western blot below. The Ponceau-red stained membrane served as the control.

Two EIF4E5 partners were already known to increase expression in the tethering assay in bloodstream forms. We found that full-length EIF4E5 also can also modestly enhance expression (Figure 2E). The combined results are consistent with a role for EIF4E5 in translation initiation in procyclic forms.

### Loss of EIF4E6 causes bloodstream-form trypanosome death

Depletion of EIF4E6 in procyclic forms had no effect on their growth, but resulted in impaired flagellar attachment (Freire *et al*., 2014a). In contrast, EIF4E6 depletion in bloodstream forms rapidly impaired growth (Figure 3A), although depletion was incomplete (Figure 3B). After 24 h, a mild decrease in protein synthesis was seen, which was more accentuated after 48h (Figure 3C), but it was difficult to tell whether this was a direct effect or a side-effect of growth inhibition.

**Figure 3.**
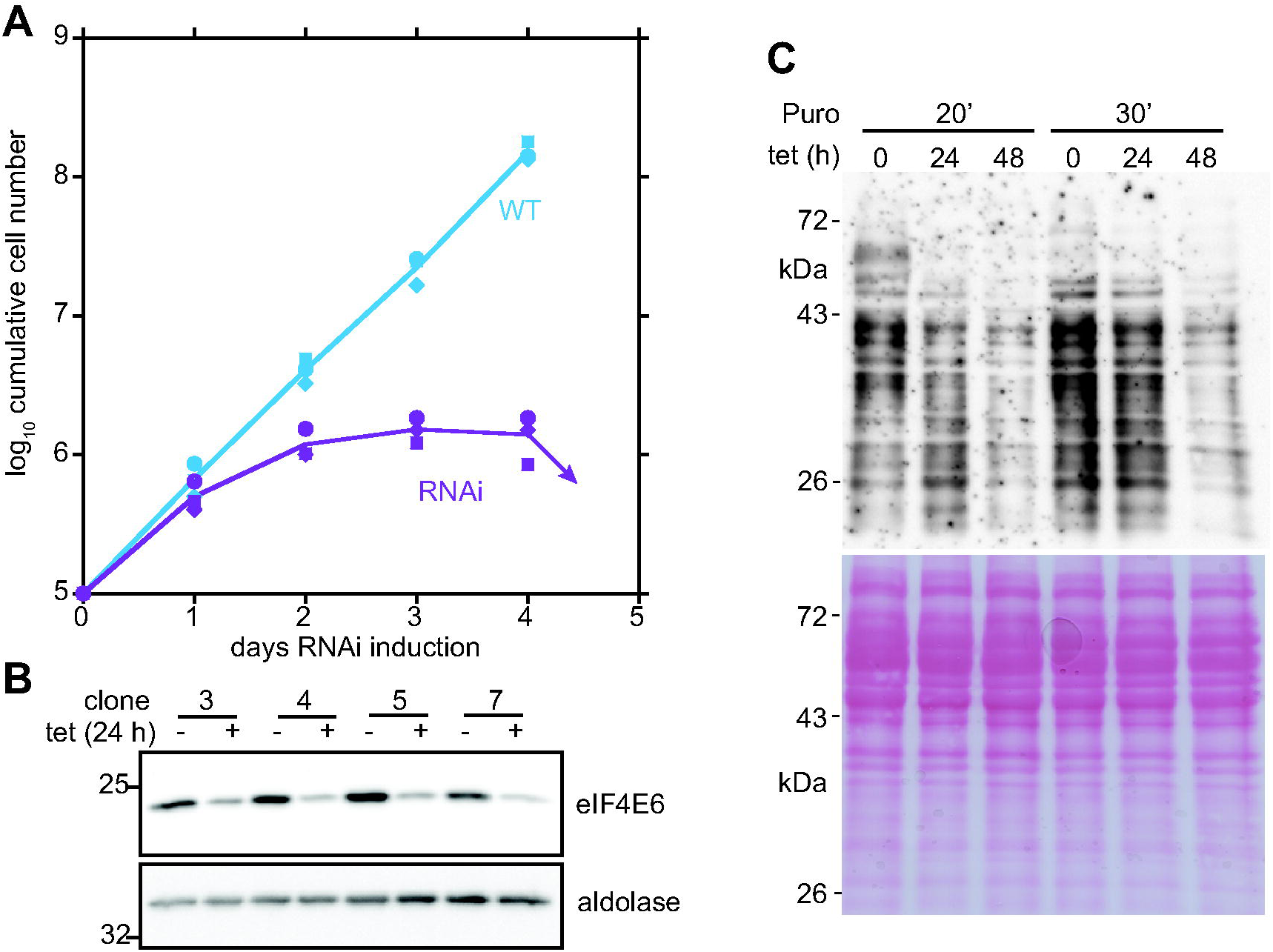
EIF4E6 is essential for bloodstream-form trypanosome growth. The level of EIF4E6 was reduced by tetracycline-inducible RNAi. **A**. Cumulative cell numbers for three different clones grown with or without 100 ng/ml tetracycline. The lines indicate the arithmetic mean values. **B**. Reduction of EIF4E6 after 24 h for four clones. EIF4E6-PTP was detected by Western blot, with aldolase as the control. The growth curves for clones 3,4 and 5 are shown in panel A. **C**. Effect of EIF4E6 depletion on translation. RNAi was induced for the times shown (no induction, 24 h or 48h). Puromycin was added for either 20 min (left) or 30 min (right) and detected by Western blotting.

To find out whether EIF4E6 is important in translation of specific mRNAs, we induced RNAi for 20h (Supplementary Figure S1A), harvesting the cells by centrifugation at maximal densities of 5 × 105/ml to ensure ongoing translation. We then purified polysomes on sucrose gradients. Four fractions were analyzed: heavy polysomes (>3 ribosomes), light polysomes (2-3 ribosomes), monosomes and large subunits, and lighter fractions. Cells with no induction served as controls. The proportions of ribosomes in the polysomal fraction were low in all samples, and hybridization with a spliced leader probe gave signals that were too weak for reliable quantification, so we could not measure the effect of the RNAi on the total polysomal distribution of mRNA. This was presumably because we added cycloheximide only after centrifuging the cells. RNASeq (shotgun sequencing of cDNA, using rRNA-depleted RNA) (Supplementary Table S3) nevertheless revealed clear differences between the polysomal fractions (light and heavy), the monosome/disome fraction, and the soluble fractions. However, there was no effect of RNAi induction (Supplementary Figure S1B). Examination of mRNAs in individual fractions with and without RNAi revealed few differences, and no signs of mRNAs that increased in one fraction while decreasing in another.

*VSG* mRNA distribution was unaffected. We thought that mRNAs with reduced translation might have lower abundance but here too there was essentially no difference with and without RNAi (Supplementary Figure S1C): the only interesting observation was that, of the five unique mRNAs that were reduced at least 1.3-fold in total mRNA after RNAi, four encoded enzymes of glucose and glycerol metabolism: glycerol kinase, glucose-6-phosphate isomerase, glyceraldehyde-3-phosphate dehydrogenase, and hexokinase (Supplementary Table S3, sheet 1). These changes may reflect the onset of growth inhibition.

### Bloodstream-form trypanosomes lacking EIF4E2 grow poorly but differentiate readily to procyclic forms

RNAi targeting EIF4E2 had previously been shown to have no effect in either bloodstream or procyclic forms, although depletion was incomplete (Freire *et al*., 2011). We were indeed able to obtain bloodstream forms that completely lacked EIF4E2 (EIF4E2 knockout, KO) (Figure 4A). However, these multiplied much slower than wild-type cells (Figure 4B) and had a maximum cell density of 8 × 10^5^/ml, in contrast to the wild-type cells, which can attain 2-2.5 × 10^6^ /ml. In contrast, after differentiation to procyclic forms, the cells without EIF4E2 grew normally (Figure 4C). We therefore decided to investigate the characteristics of EIF4E2 KO cells in more detail. First, we wondered whether the low maximum density of the cells indicated premature expression of the stumpy-form marker PAD1. Indeed, EIF4E2 KO cells already expressed PAD1 at densities of 8 × 10^5^/ml, whereas wild-type cells require a density that is four times higher (Figure 4D). To be certain that this was not an artefact of the extensive culture and cloning of the EIF4E2 KO cells, we transfected one clone with a plasmid for inducible expression of EIF4E2 (4E2 add-back). This reverted both the growth and PAD1 expression to normal levels (Figure 4D). RNAi targeting SLBP2 also decreased the growth rate (Figure 4E) and enhanced PAD1 expression (Figure 4F).

**Figure 4.**
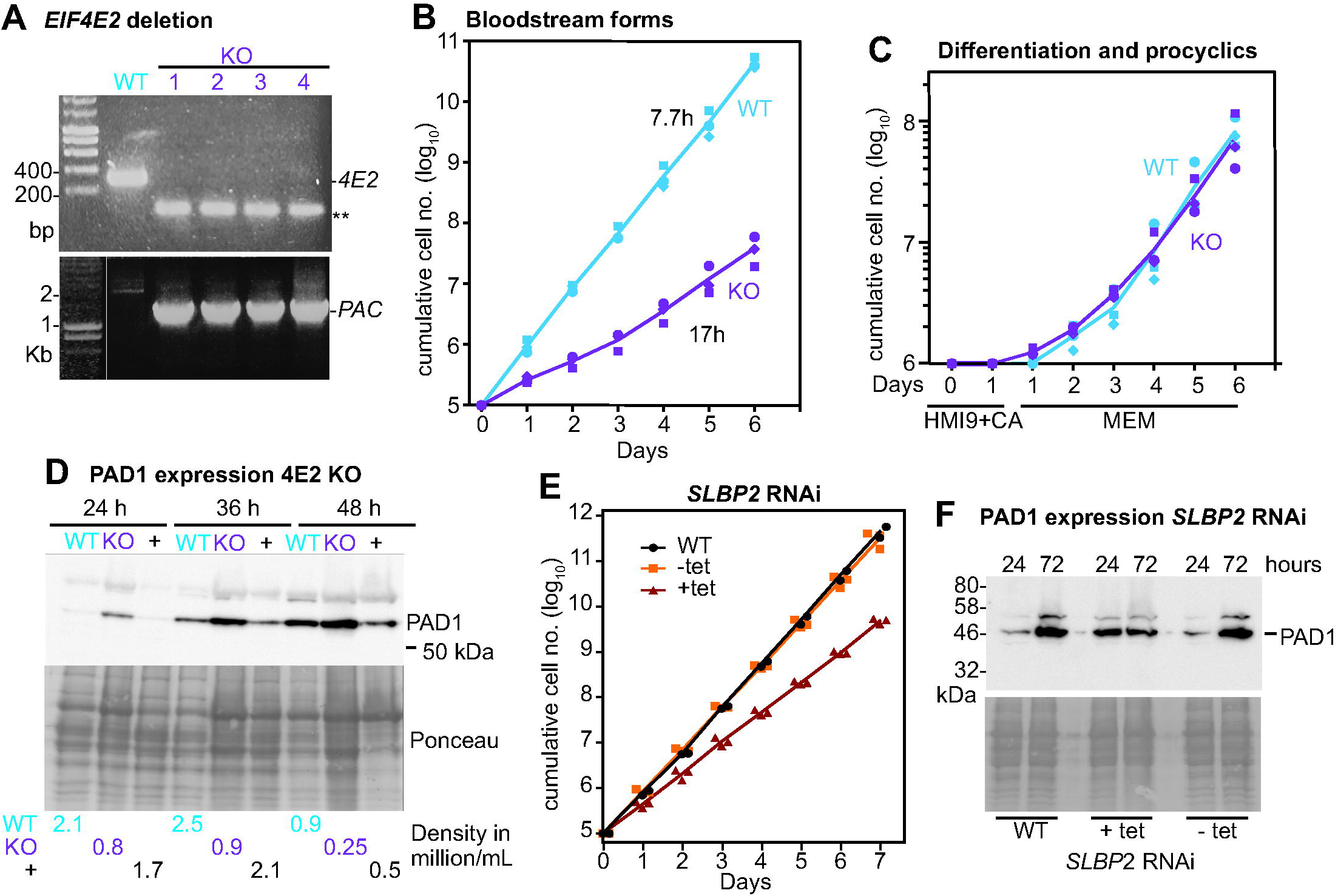
Bloodstream-form *T. brucei* lacking EIF4E2 have a growth defect. **A**. Both *EIF4E2* open reading frames were replaced by selectable markers. The *EIF4E2* gene was detected from genomic DNA using internal primers. As a control the *PAC* gene, which replaced one *EIF4E2* ORF, was amplified using a *PAC* forward primer and a reverse primer from the EIF4E2 3’-UTR. WT-wild-type; KO: knock-out. **B**. Bloodstream-form *T. brucei* lacking EIF4E2 grow slowly. Cumulative cell numbers are shown for three different clones, along with the division times. **C**. Procyclic-form *T. brucei* lacking EIF4E2 grow normally. Cells were incubated for 1 day with *cis*-aconitate at 27 °C then shifted to procyclic medium at 27 °C. Cumulative cell numbers are shown. **D**. Bloodstream-form *T. brucei* lacking EIF4E2 express PAD1 at lower densities than wildtype cells. Cells were allowed to grow to maximum density and not diluted. PAD1 is shown on the Western blot with the Ponceau-red stained membrane as control. “+” = KO cells with an inducible copy of EIF4E2. The cell densities are indicated below the lanes. **E**. Bloodstream-form *T. brucei* with reduced SLBP2 grow slowly. Cumulative cell numbers are shown for three different clones, and 2 replicates for the original cell line. **F**. PAD1 expression is induced after SLBP2 depletion.

Differentiation to stumpy forms is a quorum sensing response that is triggered by small peptides in the medium (Rojas *et al*., 2018), and can therefore be triggered in log-phase parasites by addition of medium from dense cultures (Reuner *et al*., 1997). We therefore wondered whether cells lacking EIF4E2 either (a) were excessively responsive to the differentiation signal or (b) excreted it in higher-than-normal amounts. The results of preliminary experiments with culture supernatants, however, suggested that the quorum sensing capabilities of cells without EIF4E2 were normal (Supplementary Figure S2).

### EIF4E2 specifically binds to S-phase and other mRNAs

We next searched for specific functions of the EIF4Es in bloodstream forms by cataloguing their associated mRNAs. Since cells lacking EIF4E5 grew normally, we considered only EIF4E2 and EIF4E6, again comparing with EIF4E3. The results with EIF4E3 and EIF4E6 suggested that we were detecting mainly non-specific associations. Indeed, the binding of EIF4E3 to mRNAs in bloodstream forms correlated suspiciously well with the binding of EIF4E1 to mRNAs in procyclic forms (Falk *et al*., 2021) (Figure 5A; Supplementary Table S3). The only exceptions to this were the specifically EIF4E1-associated mRNAs encoding histone H3 and NOT1, and the mRNA encoding 4EIP, which may be associated with EIF4E1 mRNA because of co-translational protein complex formation (Falk *et al*., 2021). Moreover, hardly any mRNAs were more than 4-fold enriched with either EIF4E3 or EIF4E6, and there was a moderate correlation between EIF4E3 mRNA binding and mRNA length (Figure 5B). The impression of non-specificity was accentuated when we saw the even better correlation between the binding of mRNAs to EIF4E3 and to EIF4E6 (Figure 5C; Supplementary Table S2). There was an inverse correlation between EIF4E3 mRNA association and codon adaptation score (de Freitas Nascimento *et al*., 2018), but it was very weak (Supplementary Figure S3A). There was no correlation between mRNA binding of EIF4E6 and the mRNA associations of MKT1 (Nascimento *et al*., 2020) (Supplementary Figure S3B) (Nascimento *et al*., 2020). Notably, however, EIF4E3 specifically and strongly pulled down the mRNA encoding its partner EIF4G4, while EIF4E6 did the same for its partner EIF4G5. These results suggest that EIF4G4 nascent polypeptides interact with EIF4E3, and EIF4G5 nascent polypeptides interact with EIF4E6: in other words, that the complex forms during protein synthesis (and perhaps independently of cap binding). We had previously found that EIF4E6 was associated with *VSG* mRNA, but was not detected with tubulin (*TUB*) mRNA (Melo do Nascimento *et al*., 2021). Scrutiny of our RNASeq results indeed suggested that EIF4E6 was more strongly associated with *VSG* mRNA than with *TUB* mRNA, whereas for EIF4E3, there was little difference (Figure 5C, D).

**Figure 5.**
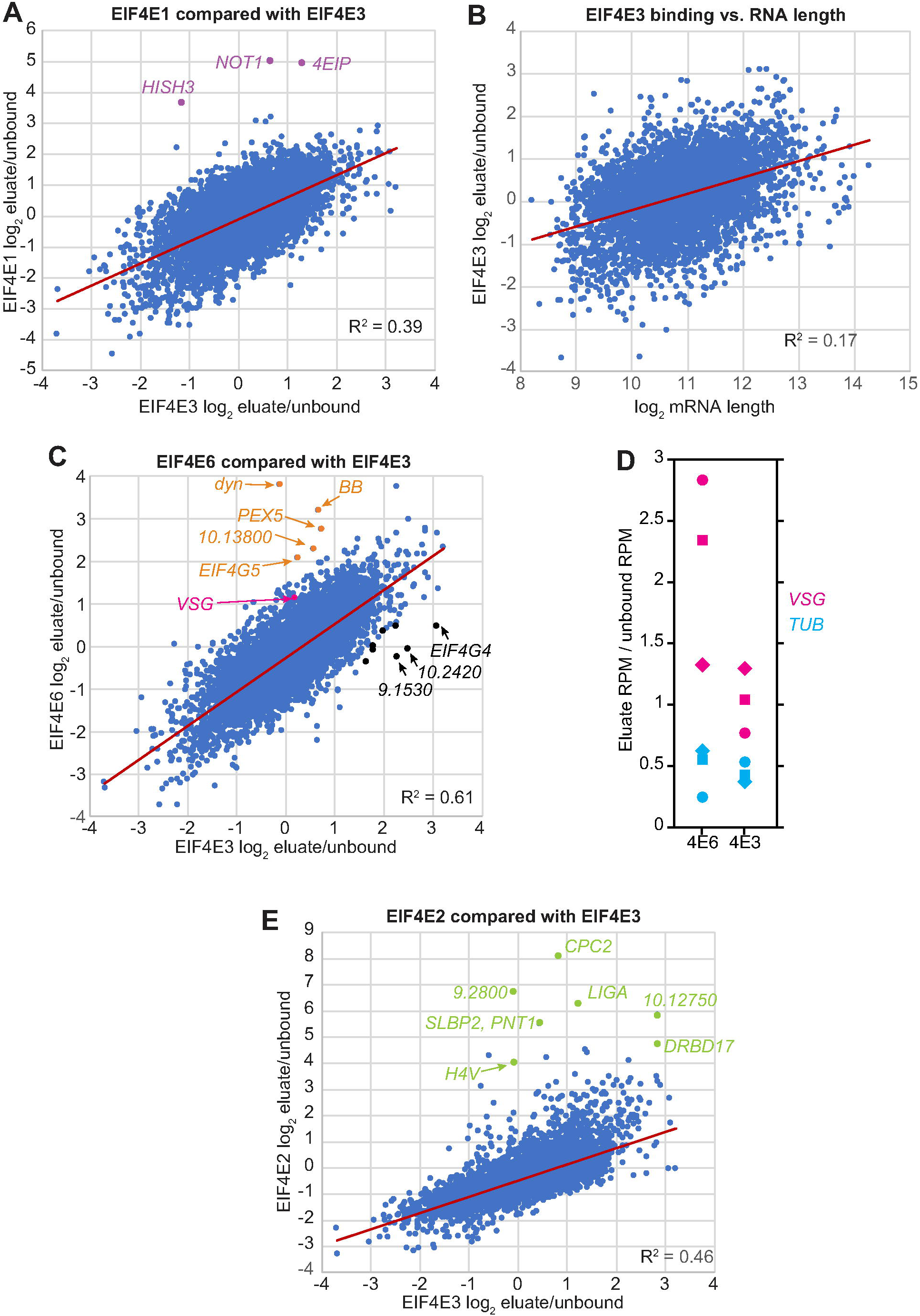
RNA binding of 4E3, 5 and 6. EIF4E-PTPs were bound to an IgG column and eluted with TEV protease. The bound RNAs were sequenced. Details are in Supplementary Table S2. **A**. Comparison of mRNAs associated with EIF4E1-PTP in procyclic forms (Falk *et al*., 2021) with those bound to EIF4E3 in bloodstream forms. For EIF4E1-PTP we used the mean ratio of bound/unbound normalized reads for one replicate with, and one without, 4EIP(Falk *et al*., 2021). For EIF4E3 the median for three replicates is shown. Association of the *4EIP*, histone H3 (*HISH3*) and *NOT1* mRNAs is reproducible (Falk *et al*., 2021) and specific to EIF4E1. The Pearson correlation coefficient was calculated in Microsoft Excel. **B**. Association of EIF4E3 with mRNAs (median value) is only very weakly correlated with mRNA length. **C**. Association of EIF4E6 with mRNAs compared with EIF4E3 (median values). The black spots indicate mRNAs for which the bound/unbound ratio with EIF4E6 was at least 3, and for which this ratio was also 3 times higher than for EIF4E6. The orange spots indicate mRNAs for which the bound/unbound ratio with EIF4E6 was at least 3, and for which this ratio was also 3 times higher than for EIF4E3. These cut-offs are arbitrary. Gene ID numbers have been shortened by removing the initial “Tb927” dyn: a dynamin; BB: a basal body protein. **D**. Association of EIF4E3 and EIF4E6 with *VSG* and tubulin (*TUB*) mRNAs. Eluate/unbound normalised reads (reads per million reads) are shown for each replicate. **E**. Association of EIF4E2 with mRNAs compared with EIF4E3 (median values). The green spots indicate mRNAs for which the bound/unbound ratio with EIF4E6 was at least 5, and for which this ratio was also 3 times higher than for EIF4E6. Unlabelled spots represent proteins with no known function.

Next, we looked at the mRNAs associated with EIF4E2. Here, too, the weaker mRNA associations correlated with those from EIF4E3 (Figure 5E) and EIF4E6 (Supplementary Figure S3C). But there was also an extremely interesting set of outliers that were enriched with EIF4E2 only. As expected, we found the mRNA encoding the protein partner SLBP2 to be bound - presumably, as for the EIF4G mRNAs described above, due to co-translational interaction. Just 18 mRNAs were at least 5-fold enriched in all replicates (Figure 5E, Table 1 and Supplementary Table S4). Of these, a few were also enriched with other RNA-binding proteins that tend to prefer long mRNAs (Supplementary Table S4, sheet 1). However it was notable that 11 of the bound mRNAs peak in S-phase according to either RNASeq results from synchronized procyclic cells (Archer *et al*., 2011), or single-cell sequencing (Briggs *et al*., 2021). (The 11 mRNAs peaked within a period of that constituted just 8% of the “pseudotime” series (Briggs *et al*., 2021).) The remaining bound mRNAs did not show significant cell cycle regulation. Strikingly, the bound mRNAs included the four S-phase-peaking transcripts that had previously been shown to be associated with the RNA-binding protein PUF9 (Archer *et al*., 2009): they encode chromosome passenger complex 2 (CPC2, Tb927.11.14840), a mitochondrial DNA ligase (lig k-alpha, Tb927.7.610), a mitochondrial protein of unknown function (PNT1, Tb927.11.6550) and a variant of histone H4 (Tb927.2.2670) (Table 1). Usually, when we examine enrichment of mRNAs with RNA-binding proteins, ratios that exceed 10 are rare. However, for EIF4E2, four ratios exceeded 40-fold in all three replicates, including those for three of the PUF9 targets (Figure 5E, Table 1 and Supplementary Table S2). Since the experiments to find PUF9-associated mRNAs were done using microarrays, it is likely that some targets were missed. PUF9 binds to mRNAs via a consensus UUGUACC motif, which was present (sometimes with a terminal mis-match) in all of the mRNAs that were at least 10-fold enriched with EIF4E2 (Table 1).

**Table 1.**
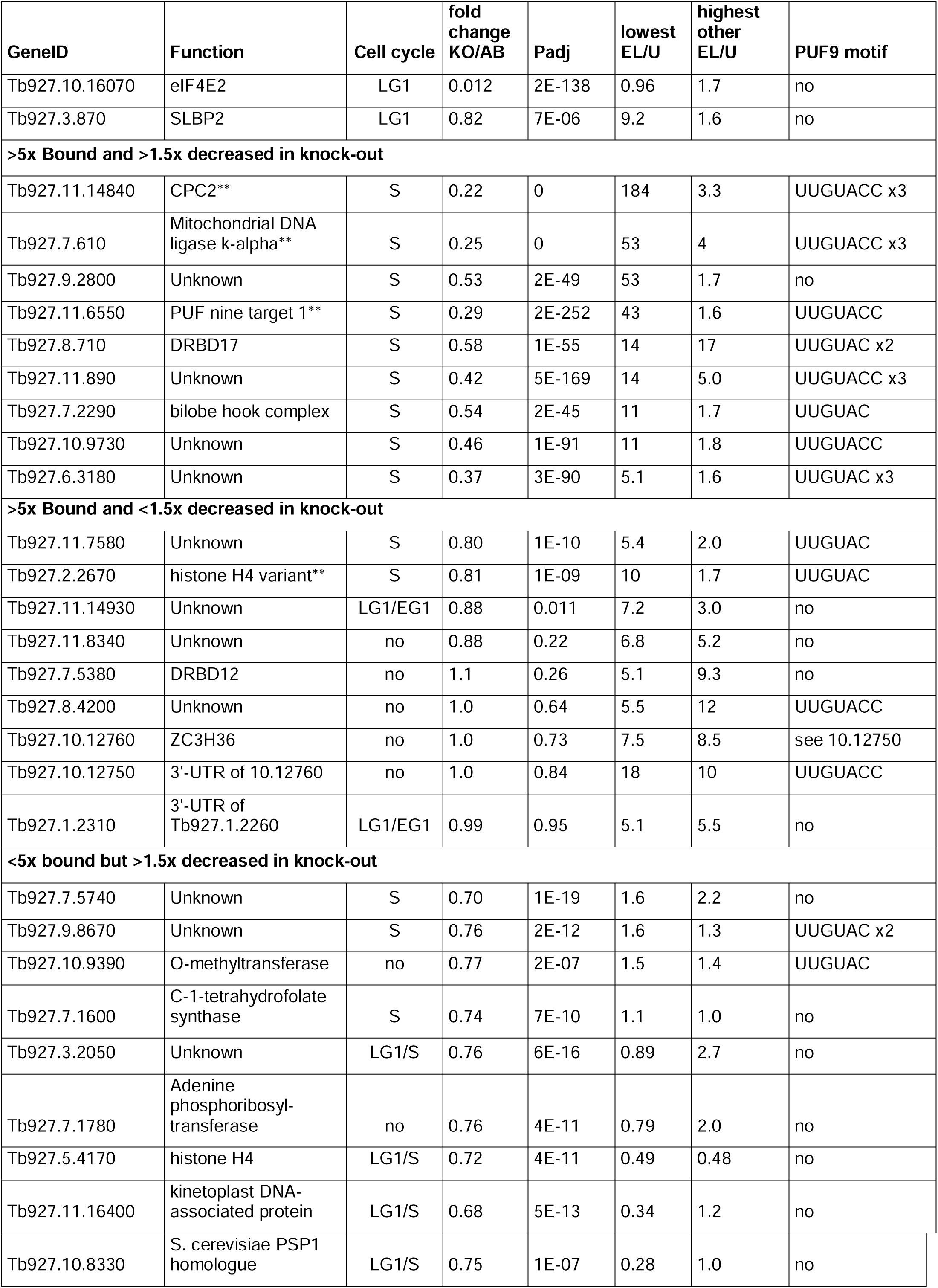
mRNAs associated with EIF4E2 The list shows all mRNAs that were either at least 5-fold enriched with EIF4E2 in all three replicates (Supplementary table S4), or showed a 1.5-fold decrease (Padj<0.05) in the EIF4E2 KO (Supplementary table S5). The top two rows are the complex components EIF4E2 and SLBP2. “Function” indicates the annotated function of the protein. **indicates mRNAs that were previously shown to bind to PUF9. “Cell cycle” shows the position of the peak mRNA amount in the cell cycle, deduced by including information from (Archer *et al*., 2011) and (Briggs *et al*., 2021). “fold change KO/AB” is the ratio of normalized reads in the KO to those in the add-back cells, and “Padj” is the adjusted P-value, both calculated using DeSeq2 (Love *et al*., 2014). “lowest EL/U” is the mRNA association with EIF4E2: we show the lowest of the three ratios of eluate to unbound normalised read (rounded to 2 significant figures). “Highest other EL/U” shows the highest eluate/unbound ration obtained for any of the other EIF4Es. “PUF9 motif” shows motifs found by manual analysis of 3’-UTRs.

| GeneID | Function | Cell cycle | fold change KO/AB | Padj | lowest EL/U | highest other EL/U | PUF9 motif |
| --- | --- | --- | --- | --- | --- | --- | --- |
| Tb927.10.16070 | eIF4E2 | LG1 | 0.012 | 2E-138 | 0.96 | 1.7 | no |
| Tb927.3.870 | SLBP2 | LG1 | 0.82 | 7E-06 | 9.2 | 1.6 | no |
| **>5x Bound and >1.5x decreased in knock-out** |  |  |  |  |  |  |  |
| Tb927.11.14840 | CPC2** | S | 0.22 | 0 | 184 | 3.3 | UUGUACC x3 |
| Tb927.7.610 | Mitochondrial DNA ligase k-alpha** | S | 0.25 | 0 | 53 | 4 | UUGUACC x3 |
| Tb927.9.2800 | Unknown | S | 0.53 | 2E-49 | 53 | 1.7 | no |
| Tb927.11.6550 | PUF nine target 1** | S | 0.29 | 2E-252 | 43 | 1.6 | UUGUACC |
| Tb927.8.710 | DRBD17 | S | 0.58 | 1E-55 | 14 | 17 | UUGUAC x2 |
| Tb927.11.890 | Unknown | S | 0.42 | 5E-169 | 14 | 5.0 | UUGUACC x3 |
| Tb927.7.2290 | bilobe hook complex | S | 0.54 | 2E-45 | 11 | 1.7 | UUGUAC |
| Tb927.10.9730 | Unknown | S | 0.46 | 1E-91 | 11 | 1.8 | UUGUACC |
| Tb927.6.3180 | Unknown | S | 0.37 | 3E-90 | 5.1 | 1.6 | UUGUAC x3 |
| **>5x Bound and <1.5x decreased in knock-out** |  |  |  |  |  |  |  |
| Tb927.11.7580 | Unknown | S | 0.80 | 1E-10 | 5.4 | 2.0 | UUGUAC |
| Tb927.2.2670 | histone H4 variant** | S | 0.81 | 1E-09 | 10 | 1.7 | UUGUAC |
| Tb927.11.14930 | Unknown | LG1/EG1 | 0.88 | 0.011 | 7.2 | 3.0 | no |
| Tb927.11.8340 | Unknown | no | 0.88 | 0.22 | 6.8 | 5.2 | no |
| Tb927.7.5380 | DRBD12 | no | 1.1 | 0.26 | 5.1 | 9.3 | no |
| Tb927.8.4200 | Unknown | no | 1.0 | 0.64 | 5.5 | 12 | UUGUACC |
| Tb927.10.12760 | ZC3H36 | no | 1.0 | 0.73 | 7.5 | 8.5 | see 10.12750 |
| Tb927.10.12750 | 3'-UTR of 10.12760 | no | 1.0 | 0.84 | 18 | 10 | UUGUACC |
| Tb927.1.2310 | 3'-UTR of Tb927.1.2260 | LG1/EG1 | 0.99 | 0.95 | 5.1 | 5.5 | no |
| **<5x bound but >1.5x decreased in knock-out** |  |  |  |  |  |  |  |
| Tb927.7.5740 | Unknown | S | 0.70 | 1E-19 | 1.6 | 2.2 | no |
| Tb927.9.8670 | Unknown | S | 0.76 | 2E-12 | 1.6 | 1.3 | UUGUAC x2 |
| Tb927.10.9390 | O-methyltransferase | no | 0.77 | 2E-07 | 1.5 | 1.4 | UUGUAC |
| Tb927.7.1600 | C-1-tetrahydrofolate synthase | S | 0.74 | 7E-10 | 1.1 | 1.0 | no |
| Tb927.3.2050 | Unknown | LG1/S | 0.76 | 6E-16 | 0.89 | 2.7 | no |
| Tb927.7.1780 | Adenine phosphoribosyl-transferase | no | 0.76 | 4E-11 | 0.79 | 2.0 | no |
| Tb927.5.4170 | histone H4 | LG1/S | 0.72 | 4E-11 | 0.49 | 0.48 | no |
| Tb927.11.16400 | kinetoplast DNA-associated protein | LG1/S | 0.68 | 5E-13 | 0.34 | 1.2 | no |
| Tb927.10.8330 | S. cerevisiae PSP1 homologue | LG1/S | 0.75 | 1E-07 | 0.28 | 1.0 | no |

### Bloodstream-form trypanosomes lacking EIF4E2 show highly specific decreases in S-phase mRNAs that are regulated by PUF9

PUF9 stabilizes its bound mRNA targets. We therefore asked whether loss of EIF4E2 would also cause a reduction in its bound mRNAs, by comparing the transcriptomes of log-phase *EIF4E2* KO trypanosomes (density ∼2 × 10^5^/ml) with those of add-back cells at the same density. There were few large changes, except for the expected decrease in *EIF4E2* mRNA itself. Only eighteen mRNAs were at least 1.5-fold significantly (Padj<0.05) decreased, but remarkably, half of them were also at least 5x enriched with EIF4E2 (Fisher P-value 2E-20). These nine mRNAs were also most strongly reduced relative to the add-back control. Of the nine decreased mRNAs that clearly were not associated with EIF4E2, eight were shown to be cell-cycle regulated by single-cell sequencing, with peaks at similar times to those of PUF9 targets. Of the remaining nine mRNAs that were bound by EIF4E2, the PUF9 target mRNA encoding histone H4V, and the mRNA from Tb927.11.7580, were only 1.2-fold, but very significantly, decreased. The remaining seven mRNAs may be unaffected by EIF4E2 deletion because they are stabilized by other interactions. We attempted to use reporters to find out whether the PUF9 binding motif was required to give poor expression in the *EIF4E2* KO cells but unfortunately, the recipient cells reproducibly failed to survive transfection.

Apart from the mildly decreased mRNAs, 35 mRNAs were increased at least 1.5-fold (Padj <0.05) in the *EIF4E2* KO cells. None of these is bound by EIF4E2. Usually, each trypanosome transcribes only one *VSG* gene, from a telomeric “expression site”. Although amounts of mRNA encoding the major VSGs (EATRO1125_VSG211 and EATRO1125_VSG11) were unaffected by the EIF4E2 KO, mRNAs from the 16 other expression site *VSG* genes that we examined increased in abundance. This increase was from an extremely low base, such that less than 1 in 50 KO cells would be expected to express even one additional *VSG* mRNA. Nevertheless, the result suggests that the KO has caused slight de-regulation of VSG expression. Most of the other increased mRNAs encode proteins of unknown function, but they did include those encoding ESAG9 and the procyclic marker procyclin, which are elevated in stumpy forms (Silvester *et al*., 2018).

### The cell cycle of trypanosomes lacking EIF4E2

Our transcriptomic results led us to suspect that EIF4E2 and SLBP2 collaborate with PUF9 to stabilise specific mRNAs at the start of S-phase. To find out whether the cells lacking EIF4E2 have a cell cycle defect, we first counted nucleus and kinetoplasts. The kinetoplast is a DNA structure consisting of multiple concatenated copies of mitochondrial DNAs - “maxicircles” which contain the protein-coding and rRNA genes, and “minicircles” which are needed for mitochondrial mRNA editing. Cells with one nucleus are in G1 or S, those with 1 nucleus and 2 kinetoplasts are in G2 or M, and the presence of 2 nuclei indicates completed mitosis prior to cytokinesis. Counting of stained smears (at least 100 cells) revealed no significant differences between the KO cells and the add-back. However, this method does not allow us to detect S-phase or discrepancies in nuclear DNA content. We therefore also analysed the DNA content of the cells by propidium iodide staining and FACScan. This revealed a slight enrichment of cells with a 4N DNA content (G2) at the expense of G1 (Figure 6A).

**Figure 6.**
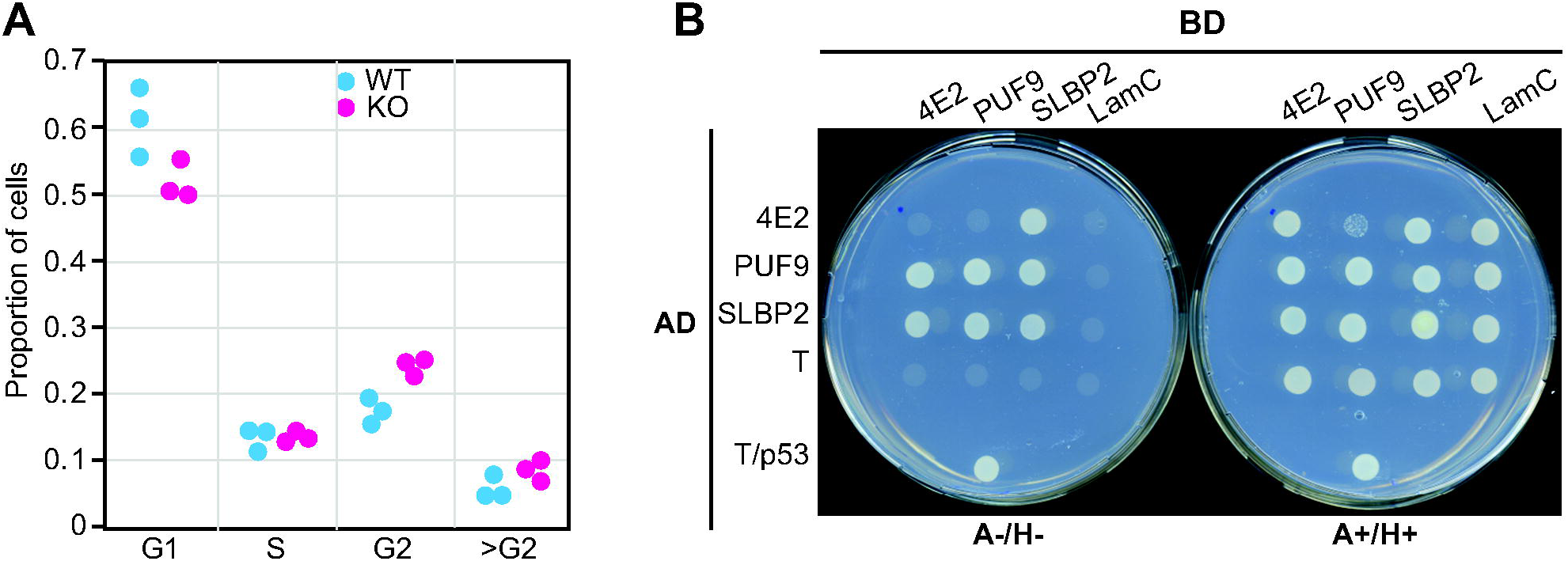
Functions of EIF4E2, SLBP2 and PUF9. **A**. Wild-type cells or cells lacking EIF4E2 were fixed, stained with propidium iodide, and analysed by FACScan. The cells were then categorized according to DNA content (2N= G1, >2n and <4N S, 4N = G2, and >4N (abnormal). **B**. EIF4E2, SLBP2 and PUF9 were expressed as fusions with the DNA-binding domain or the transcription activating domain of the GAL4 transcription factor, in a suitable yeast reporter strain. Images of two different plates are shown. The right-hand plate selects for both plasmids but contains adenine and histidine. The left-hand plate lacks adenine and histidine, and cells will only grow if the DNA-binding-domain and activation-domain fusion proteins interact. Similar results were obtained in three experiments.

### EIF4E2 and SLBP2 can interact with PUF9

Our mass spectrometry results for purified EIF4E2-PTP revealed copurification only of SLBP2 (Supplementary Table S2). PUF9 was detected in only one of the five replicates, but this sample also had a large degree of contamination with other proteins. Our results therefore indicate that only a small proportion of - if any - EIF4E2 is associated with PUF9 *in vivo*. To find out whether either EIF4E2 or SLBP2 can even interact with PUF9, we used a yeast 2-hybrid assay. Results (Figure 6B) revealed a clear interaction between EIF4E2 and SLBP2. There was also interaction between PUF9 and EIF4E2 when EIF4E2 was fused to the DNA-binding domain and PUF9 was fused with the activation domain, but for the opposite orientation, co-expression was reproducibly and toxic so no firm conclusions could be drawn. PUF9 also interacted with itself. These results suggest that SLBP2 can interact with PUF9 and that the interaction between SLBP2 and PUF9 does not require a trypanosome-specific protein modification.

## Discussion

We here investigated the roles of EIF4E2, EIF4E5, and EIF4E6 in bloodstream-form *T. brucei*, by identifying protein interaction partners and bound mRNAs, and by assessing the effects of depletion. This yielded few novel insights for EIF4E5 and EIF4E6, but an unexpected role for EIF4E2 in stabilizing S-phase mRNAs.

EIF4E5 co-purified the same proteins in bloodstream forms as in procyclic forms, with PABP2 but not PABP1. EIF4E5 was dispensable for growth of bloodstream forms *in vitro*, but cells that had differentiated to the procyclic form failed to multiply; procyclic forms in which EIF4E5 was only depleted were already known to have a motility defect. Thus so far, the functions of EIF4E5 appear to be restricted to forms that multiply in the vector, although examination of the behavior of cells lacking EIF4E5 during mammalian infection might reveal roles that were not seen *in vitro*.

For EIF4E6, we confirmed preferential association with PABP2, MKT1 and its partner PBP1, as well as CFB2 and the *VSG* mRNA. Our studies of EIF4E6 also revealed there were also a few additional proteins, perhaps associated with bound mRNAs. For example, ZFP2 and DRBD2 (Wippel *et al*., 2019) are cytosolic proteins; DRBD2 was reported to associate with PABP2 (Assis *et al*., 2021, Kramer *et al*., 2013), and ZFP2 is required for differentiation of bloodstream forms the procyclic form (Hendriks *et al*., 2001). DRBD3, which was present with EIF4E6 but more with EIF4E3, probably influences splicing but can also stabilize bound mRNAs (Das *et al*., 2015, Fernandez-Moya *et al*., 2012, Stern *et al*., 2009, Estévez, 2008). TRRM1 is primarily in the nucleus (Naguleswaran *et al*., 2015). EIF4E6 depletion inhibited trypanosome proliferation and protein synthesis, (although we did not demonstrate that the effect on protein synthesis was direct). EIF4E6 and its partner EIF4G5 are as abundant as EIF4E3 and EIF4G4, and both complexes are associated with several of the same RNA-binding proteins. We therefore suggest that trypanosome EIF4E6 is an active translation factor, especially for mRNAs bound by MKT1, but perhaps with a function that overlaps with that of EIF4E3. We do not know why over-expression of EIF4E6 in *Leishmania* inhibited translation: one possibility is that EIF4E6 is indeed a translational repressor, but another possibility is a “squelching”-type phenomenon. In the over-expressing cells, the *EIF4E6* mRNA was at least 200-fold more abundant than normal. If EIF4E6 protein was also very highly over-expressed, this might have enabled it to compete with EIF4E4 or EIF4E3 for binding to *Leishmania* EIF4G3 or EIF4G4.

It was notable that for each EIF4E that we examined, the mRNA encoding the known direct binding partner protein was pulled down: *4EIP* mRNA with EIF4E1 (Falk *et al*., 2021); *SLBP2* mRNA with EIF4E2; *EIF4G4* mRNA with EIF4E3; and *EIF4G5* mRNA with EIF4E6. We suggest that in each of these cases, the nascent polypeptide of the interacting protein binds to its partner. There is extensive evidence for this phenomenon in other organisms (Duncan & Mata, 2011, Shiber *et al*., 2018, Panasenko *et al*., 2019, Lautier *et al*., 2021). Stabilization of each protein by the other can also explain why the partners are in most cases present at similar abundances (Supplementary Table S1). Apart from this, with the exception of RNAs bound specifically by EIF4E2, and two mRNAs bound by EIF4E1 (Falk *et al*., 2021), the results of the RNA pull-downs for all of EIF4Es were suspiciously similar, suggesting that we were detecting mainly non-specific interactions - that is, that the proteins are mostly only transiently associated with target mRNAs. After recruitment of the 43S complex, there are two possible fates for associated EIF4F. In some mammalian cell types, EIF4F usually stays bound to the 43S complex until 60S joining occurs (cap-tethered scanning), so that only one 43S complex can scan at a time (Bohlen *et al*., 2020). This results in less efficient translation of mRNAs with long 5’-UTRs (Bohlen *et al*., 2020). In contrast, in other mammalian cell types and yeast, the association is lost after the 43S complex scans the 5’-UTR, so that more than one 43S complex can be present on a 5’-UTR. In that case, translation efficiency is not decreased by long 5’-UTRs (unless the 5’-UTR contains an open reading frame) (Archer *et al*., 2016, Shirokikh *et al*., 2017). Ribosome densities on trypanosome mRNAs (Antwi *et al*., 2016) are not affected by long 5’-UTRs (Supplementary Figure S4), suggesting that as in yeast, scanning is not cap-tethered. This means that in principle, EIF4F could be lost as soon as scanning has started.

EIF4E2 was associated with mRNAs that are also bound by PUF9. Moreover, the abundances of these mRNAs were lower in the absence of EIF4E2 than in its presence, suggesting that EIF4E2 stabilizes PUF9-bound mRNAs. Two-hybrid assay results also suggested an interaction between PUF9 and SLBP2, and possibly also EIF4E2, but PUF9 was not detected with purified EIF4E2-PTP, even though PUF9 protein is (according to mass spectrometry results) at least twice as abundant as SLBP2 and EIF4E2 (Dejung *et al*., 2016, Benz & Urbaniak, 2019, Crozier *et al*., 2018, Tinti & Ferguson, 2022). Although PUF9 is larger than SLBP2 and EIF4E2, and many factors can affect mass spectrometry detection, this suggests that EIF4E2 and SLBP2 are not present in massive excess. We therefore suggest that the EIF4E2-SLBP2 complex and PUF9 interact only briefly, at a specific stage of the cell cycle. The *EIF4E2* and *SLBP2* mRNAs both peak in late G1 (Archer *et al*., 2011). PUF9 has low abundance in G1, beginning to increase in early S-phase and also showing increased phosphorylation at serine 617, which is close to a poly(Q) region (Benz & Urbaniak, 2019). There is also a 30-fold increase in phosphorylation at threonine 400 of SLBP2 between early G1 and S-phase (Benz & Urbaniak, 2019). It is also possible that an *in vivo* interaction between the EIF4E2/SLBP2 complex and PUF9 is stabilised by cooperativity, via sequence-specific binding of PUF9 to its target mRNAs together with EIF4E2 binding to the cap.

EIF4E2 target mRNAs are mostly cell-cycle regulated, with a peak during S-phase - after accumulation of PUF9 and the mRNAs encoding EIF4E2 and SLBP2. There are two possibilities for EIF4E2/SLBP2/PUF9 function. One possibility is that the complex is able to act in translation initiation. Although absolutely no translation initiation factors were detected in the EIF4E2 purifications, low-stoichiometry interactions specific to a particular cell-cycle stage (when PUF9 is also associated) would not have been detected. The alternative is that the EIF4E2/SLBP2/PUF9 complex acts to store accumulated RNAs in a non-translated state, before releasing them to enable specifically-timed translation initiation via a different EIF4E complex - perhaps as a consequence of the phosphorylations noted above. Unfortunately, we do not know when most of the proteins encoded by the target mRNAs increase in abundance, since they were not detected in cell-cycle proteome studies (Crozier *et al*., 2018, Benz & Urbaniak, 2019) and ribosome profiling results are not available for synchronized cells.

EIF4E2 depletion caused a very slight increase in G2 cells, which is presumably caused by decreases in the amounts of proteins encoded by its target mRNAs. The proteins have diverse functions, and decreases might be expected to cause defects in division of both the nucleus and the kinetoplast, without affecting DNA synthesis. The bound mRNA that was most strongly affected by EIF4E2 (or PUF9) depletion encodes CPC2 (Tb927.11.14840), a kinetoplastid-specific component of the chromosome passenger complex. Chromosome passenger complexes are required for spindle assembly and chromosome segregation, and depletion of CPC2 causes an increase in cells in G2, as judged by their having 4N DNA content and 2 kinetoplasts, but only one nucleus (Li *et al*., 2008). The next most strongly affected mRNA (Tb927.11.6550) encodes a mitochondrial protein, PNT1 (PUF9 target 1). Expression of C-terminally tagged PNT1 in addition to the wild-type caused accumulation of extra kinetoplasts, while RNAi inhibited trypanosome division (Archer *et al*., 2009). Both of these mRNAs were also pulled down with PUF9, as was that encoding the mitochondrial DNA ligase k-alpha (Tb927.7.610) (Archer *et al*., 2009). Depletion of DNA ligase k-alpha prevents re-ligation of minicircle DNAs, a major component of the kinetoplast DNA assemblage, and inhibits kinetoplast segregation, ultimately resulting in cells that lack the kinetoplast (Downey *et al*., 2005). The other target mRNA with a known function encodes histone H4v, which is located at sites of RNA polymerase II transcription termination (Siegel *et al*., 2009). The targets also included two mitochondrial/kinetoplast-associated proteins of unknown function (Tb927.11.890, Tb927.11.7580), and one (Tb927.7.2290) that is located in the bilobe, a structure located near the basal body (Dean *et al*., 2016, de Graffenried *et al*., 2013).

In cells without EIF4E2 we saw, relative to “add-back” cells containing EIF4E2, elevated amounts of various mRNAs that increase in stumpy-form trypanosomes. None of these mRNAs was bound by EIF4E2 so the effects are probably secondary to stress (Quintana *et al*., 2021). Another indication of stress in the cells without EIF4E2 was the detection of small amounts of numerous different *VSG* mRNAs in addition to the major one from the active expression site. This phenomenon is known to be induced by DNA damage (Black *et al*., 2020) and by loss of histone H4v (Müller *et al*., 2018). These cells showed good competence for differentiation into procyclic forms, as might be expected for cells with partially stumpy-like transcriptomes.

A major question that remains is why cells without EIF4E2 are able to grow at normal rates as procyclic forms. PUF9 depletion also did not affect procyclic-form growth (Archer *et al*., 2009). Although cell cycle regulation may be different in procyclic forms, there is also a trivial explanation: procyclic forms have a slower growth rate than bloodstream forms in culture (division times of about 11 hours in contrast to about 6 hours for bloodstream forms) so perhaps lower levels of the PUF9/EIF4E2 targets can be tolerated. The presence of EIF4E2, SLBP2 and PUF9 in *Crithidia fasciculata*, and of EIF4E2 and SLBP2 in free-living *Bodo saltans*, suggests that the functions of this complex are not restricted to parasitism of vertebrates. It would therefore be interesting to find out how procyclic forms without EIF4E2 perform in the more stressful environment of the natural host.

## Materials and methods

### Trypanosome culture and differentiation

Pleomorphic EATRO1125 ectopically expressing the *Tet* repressor were used. Bloodstream forms were cultured at 37 °C in HMI-9 medium (supplemented with 10 % (v/v) fetal calf serum (FCS), 1 % (v/v) penicillin/streptomycin solution (Labochem international, Germany), 15 μM L-cysteine, and 0.2 mM β-mercaptoethanol) in the presence of 5 % CO_2_ and 95 % humidity and their density was usually maintained below 1.0 × 10^6^ cells/ml. Procyclic forms were cultured at 27 °C in MEM-Pros media (supplemented with 10 % (v/v) FCS, 3.75 mg hemin, and 1 % (v/v) penicillin/streptomycin solution (Labochem international, Germany)), and their density was maintained between 5 × 10^5^ and 5 × 10^6^ cells/ml.

To induce bloodstream form to procyclic form differentiation, cells grown at 1 × 10^6^ cells/ml in HMI-9 medium were incubated with 6 mM *cis*-aconitate at 27 °C for 24 h. On the following day, the medium was exchanged for MEM and the cells were cultured at 27 °C afterwards.

### Cell cycle analysis

Cells were attached to slides, fixed, then labelled with DAPI at 500ng/ml (Falk *et al*., 2021). Images were acquired using an Olympus IX81microscope and analyzed with Image J software. Counting of at least 50 cells per sample was done using blinded images. Alternatively, Cells were collected by centrifugation, resuspended in 500 μl of PBS with 10μg/ml RNase A and 30 μg/ml propidium iodide, incubated at 37°C for 30 min and analysed by FACSSCAN.

### Genetic manipulation of trypanosomes

Gene knockout was achieved by replacing the open reading frame of *EIF4E2* with open reading frames encoding blasticidin and puromycin resistance genes, which integrate by homologous recombination. Double knock-out cells were selected by growing transfectants in both 5 μg/ml blasticidin and 0.2 μg/ml puromycin for at least two weeks before downstream analyses. Tagging with the PTP-tags was achieved by integration through homologous recombination at the C-terminus followed by selection of transfectants. The primers and plasmids used in this study can be found in Supplementary Table S3.

### DNA extraction and PCR

DNA was extracted from approximately 3 × 10^8^ cells as follows. The cell pellet was resuspended and the cells were lysed using 0.5 ml EB buffer (10 mM Tris-HCl pH 8.0, 10 mM NaCl, 10 mM EDTA). RNA was digested using 25 μg/ml of RNaseA (Sigma-Aldrich) at 37 °C for 30 min. Proteins were precipitated using 1.5 M ammonium acetate and centrifuging at 9,391 × *g* for 5 min. Isopropanol was added to the supernatant in a 1:1 ratio and centrifuged at 15,871 × *g* for 15 min at 4 °C. The pellet was washed with 75 % ethanol followed by centrifugation at 15,871 × *g* for 5 min and dried before resuspending in 50 μl TE buffer (10 mM Tris pH 7.5, 1 mM EDTA pH 8.0). PCR was done using Taq Polymerase according to the manufacturer’s instructions (New England Biolabs).

### Preparation of protein and RNA for mass spectrometry

Pull-downs were performed with 10^9^ bloodstream forms as previously described (Falk *et al*., 2021). Briefly, cells were harvested, washed, then stored at -80 °C until use. Thawed cells were resuspended in 0.5 ml of lysis buffer (20 mM Tris [pH 7.5], 5 mM MgCl_2_, 1 mM DTT, 0.05 % IGEPAL, 100 U/ml RNasin, 10 μg/ml aprotinin, and 10 μg/ml leupeptin), and passed 20× through a 21G×1½’’ needle and 20× through a 27G×¾ needle. Samples were then cleared at 10,000 × *g* for 15 min. The salt concentration was adjusted to 150 mM KCl. Magnetic beads (Dynabeads™ M-280 Tosylactivated, Thermo Fisher Scientific) coupled to rabbit IgG were washed three times with (20 mM Tris [pH 7.5], 5 mM MgCl_2_, 1 mM DTT, 0.05% IGEPAL, 100 U/ml RNasin, 150 mM KCl). 100 μl of the beads were then added and the mixture was incubated for 1-2h at 4 °C with rotation. The beads were washed four times with wash buffer, then, bound proteins were released by incubation for 90 min at 20 °C with TEV protease (1 mg/ml). The supernatant was transferred to a fresh tube, and 10 μl of equalization buffer (200 mM sodium phosphate, 600 mM sodium chloride, 0.1 % Tween-20, 60 mM imidazole, pH 8.5), as well as 30 μl of Ni-NTA-magnetic beads were added. After incubation (30 min at 20 °C) the supernatant was collected and stored in Laemmli buffer at - 80 °C.

Eluted proteins were separated on a 1.5 mm NuPAGE™ Novex™ 4-12 % Bis-Tris protein gel (Thermo Fisher Scientific) until the running front had migrated roughly 2 cm, after which the gel was stained with Coomassie blue and de-stained with de-staining solution (10 % acetic acid, 50 % methanol in H_2_O). Three areas per lane were cut and analyzed by Nanoflow LC-MS2 analysis with an Ultimate 3000 liquid chromatography system directly coupled to an Orbitrap Elite mass spectrometer (both Thermo-Fischer, Bremen, Germany). MS spectra (m/z 400–1600) were acquired in the Orbitrap at 60,000 (m/z 400) resolution. Fragmentation in CID cell was performed for up to 10 precursors. MS2 spectra were acquired at rapid scan rate. Raw files were processed using MaxQuant (version 1.5.3.30; J. Cox, M. Mann, Nat Biotechnol 2008, 26, 1367) for peptide identification and quantification. MS2 spectra were searched against the TriTrypDB-8.1TREU927-AnnotatedProteins-1 database (containing 11567 sequences). Data were analyzed quantitatively and plotted using Perseus software (Version 1.6.6.0). The mass spectrometry proteomics data have been deposited to the ProteomeXchange Consortium via the PRIDE partner repository with the dataset identifiers PXD025913 and PXD033511.

For the RNA-binding assays, the binding was done as above. Unbound samples were collected for RNA extractions, three volumes of peqGOLD TriFast™ FL reagent were added, and samples were stored at -80 °C until further processing. The beads were washed four times with wash buffer and proteins with bound RNAs were released by incubation with 5 μl recombinant TEV protease (1 mg/ml) in 250 μl of wash buffer for 90 min at 20 °C. For collecting the elution fractions, the beads were concentrated on one side of the tube, the supernatant was transferred to a fresh tube, three volumes of peqGOLD TriFast™ FL reagent were added, and samples were stored at -80 °C until further processing.

### RNA isolation

Polysomes were prepared on sucrose gradients as previously described (Minia *et al*., 2016, Minia & Clayton, 2016), but without cycloheximide pre-treatment. Whole cell pellets or gradient fractions were stored at -80 °C in TriFast reagent until use. For RNA preparation, the samples were thawed and incubated for 5 min at room temperature to ensure complete dissociation of ribonuclear complexes before isolation according to the manufacturer’s instructions.

### RNA sequencing and data analysis

RNA was subjected to rRNA depletion using oligonucleotides and RNase H (Minia *et al*., 2016). RNA-seq was done at the CellNetworks Deep Sequencing Core Facility at the University of Heidelberg. For library preparation, NEBNext Ultra RNA Library Prep Kit for Illumina (New England BioLabs Inc.) was used. The libraries were multiplexed (6 samples per lane) and sequenced with a Nextseq 550 system, generating 75 bp single-end sequencing reads. The data were aligned and to the TREU 927 reference genome, and to a small set of expressed EATRO1125 *VSG* coding regions, using a custom python script (Leiss *et al*., 2016) Briefly, the quality of the data was checked using FastQC, primers were removed using Cutadapt (Martin, 2011), and the data were aligned using Bowtie2 (Langmead & Salzberg, 2012). Differential expression was analyzed using a specialized version (Leiss & Clayton, 2016) of DeSeq2 (Love *et al*., 2014). To assess enrichment of particular functional classes we restricted the analysis to a list of “unique” genes containing only one representative each of repeated or similar genes (modified from (Siegel *et al*., 2010)). The data are available at the Annotare part of Array Express (https://www.ebi.ac.uk/fg/annotare) with the accession numbers E-MTAB-10670 (mRNAs associated with EIF4E2) E-MTAB-10453 (mRNAs associated with EIF4E1 and EIF4E6), E-MTAB-10665 (mRNAs associated with EIF4E3), E-MTAB-10719 (comparison of EIF4E2 knock-out and add-back) and E-MTAB-11714 (EIF4E6 RNAi).

### Protein detection by western blotting

For western blotting, 1-5 × 10^6^ cells were lysed in 1× Laemmli buffer, separated by 8-12 % SDS-PAGE, and processed as described in (Klein *et al*., 2015). The following antibodies were used for specific protein detection: anti-4E1 serum (1:2000, rabbit; kind gift from Osvaldo de Melo Neto), anti-S9 serum (1:20,000, rat; loading control), anti-myc (1:5000, mouse, Y2H), anti-p-GPEET serum (1:2000, western blotting), anti-PAP, anti-EP,

### Cell cycle and DNA content analysis

For FACScan analysis of DNA content, the parasites were washed twice with phosphate-buffered saline (PBS) and fixed in methanol for 1 h at 4 °C. The cells were washed twice again with PBS and suspended in PBS. The cells were incubated for 30 min at room temperature with DNase-free RNase (10 μg/ml) and propidium iodide (20 μg/ml) before analysis with a fluorescence-activated cell sorting-scan (FACScan) analytical flow cytometer (BD Biosciences). Percentages of cells in each phase of the cell cycle (G1, S, and G2/M) were determined using the FlowJo software.

## Acknowledgements

We thank Claudia Helbig for technical assistance, and Osvaldo de Melo Neto (Fiocruz, Recife, Brazil) for providing plasmids and for useful discussions. Sequencing was done by David Ibberson (Deep Sequencing Core Facility), and Thomas Ruppert and Sabine Merker (Core Facility for Mass Spectrometry) performed the mass spectrometry. This project was mainly supported by grant number Cl112/30 from the Deutsche Forschungsgemeinschaft.

## Author contributions

FF designed and performed most of the experiments and did the relevant data analysis. RMP was responsible for the yeast 2-hybrid analysis, cell-cycle analysis, the second series of EIF4E2-PTP preparations, and polysomal RNA preparation. AW analyzed the EIF4E2 mass spectrometry results and the EIF4E6 RNAi results. CC was responsible for conceptualization, funding acquisition, supervision, and project administration. CC wrote the manuscript, partly based on FF’s PhD thesis.

