## Supplementary for "Roles and interactions of the specialized initiation factors EIF4E2, EIF4E5 and EIF4E6 in *Trypanosoma brucei*: EIF4E2 maintains the abundances of S-phase mRNAs"

**Supplementary Figures:**

**Supplementary Figure S1. RNASeq for cells with *EIF4E6* RNAi**

**A.** Western blot showing EIF4E6-PTP depletion

**B.** Principal component analysis for all fractions, all unique coding regions (Supplementary Table S3)

**C.** Principal component analysis for total mRNA, all unique coding regions (Supplementary Table S3)

**Supplementary Figure S2. Cells lacking EIF4E2 react normally to high density supernatants**.

**A-F**: Filtered supernatants were taked from wild-type cells grown to maximum density. Wildtype (A,B), knock-out (C,D) or add-back cells (E,F) were seeded at starting densities of 1 x $10^5$ cells/ml (A,C,E) or 2.5 x $10^5$ cells/ml (B,D,F) with fresh HMI-9, maximum density supernatant (MDS), or MDS diluted 1:3 or 1:5 with fresh HMI-9. Cells were diluted after 24 h as necessary.

**G, H**: EIF4E2 KO cells, or add-back cells, were incubated for 24,30 or 36h at maximum density and filtered supernatants were prepared. Wild-type cells were then resuspended in the supernatants at 1 x $10^5$ cells/ml. The maximum densities for EIF4E2 KO cells is 8 x $10^5$ cells/ml, and for add-back cells, 2.5 x $10^6$ cells/ml.

**Supplementary Figure S3. Additional correlations for RNA binding.**

EIF4E-PTPs were bound to an IgG column and eluted with TEV protease. The bound RNAs were sequenced. Details are in Supplementary Table S2.

**A.** Comparison between EIF4E3 RNA binding (eluate/unbound) and codon optimality, as judged by the gCAI (de Freitas Nascimento *et al.*, 2018).

**B.** Comparison between EIF4E6 RNA binding (eluate/unbound) and MKT1 RNA binding (Nascimento *et al.*, 2020).

**C.** Comparison between EIF4E2 RNA binding (eluate/unbound) and EIF4E6 RNA binding.

**Supplementary Figure S4.**

Relationship between ribosome density (Antwi *et al.*, 2016) and 5'-UTR length.

**Supplementary Tables:**

All Tables have legends on the first sheet.

**Supplementary Table S1.**

Abundances of translation factors by quantitative mass spectrometry (Tinti & Ferguson, 2022)

**Supplementary Table S2.**

Mass spectrometry of purified EIF4E-PTP preparations

**Supplementary Table S3.**

RNASeq of total and sucrose-gradient-fractionated RNA after EIF4E6 depletion

**Supplementary Table S4.**

RNASeq of RNAs bound to various EIF4Es, with EIF4E1 (Falk *et al.*, 2021), 4EIP (Terrao *et al.*, 2018)and MKT1 (Nascimento *et al.*, 2020) for comparison.

**Supplementary Table S5.**

RNASeq of RNA in cells lacking EIF4E2, compared with cells having EIF4E2

**Supplementary Table S6.**

Plasmids and oligonucleotides
