## Supplementary figures and images for "Roles and interactions of the specialized initiation factors EIF4E2, EIF4E5 and EIF4E6 in *Trypanosoma brucei*: EIF4E2 maintains the abundances of S-phase mRNAs"

### Supplementary Figure S1

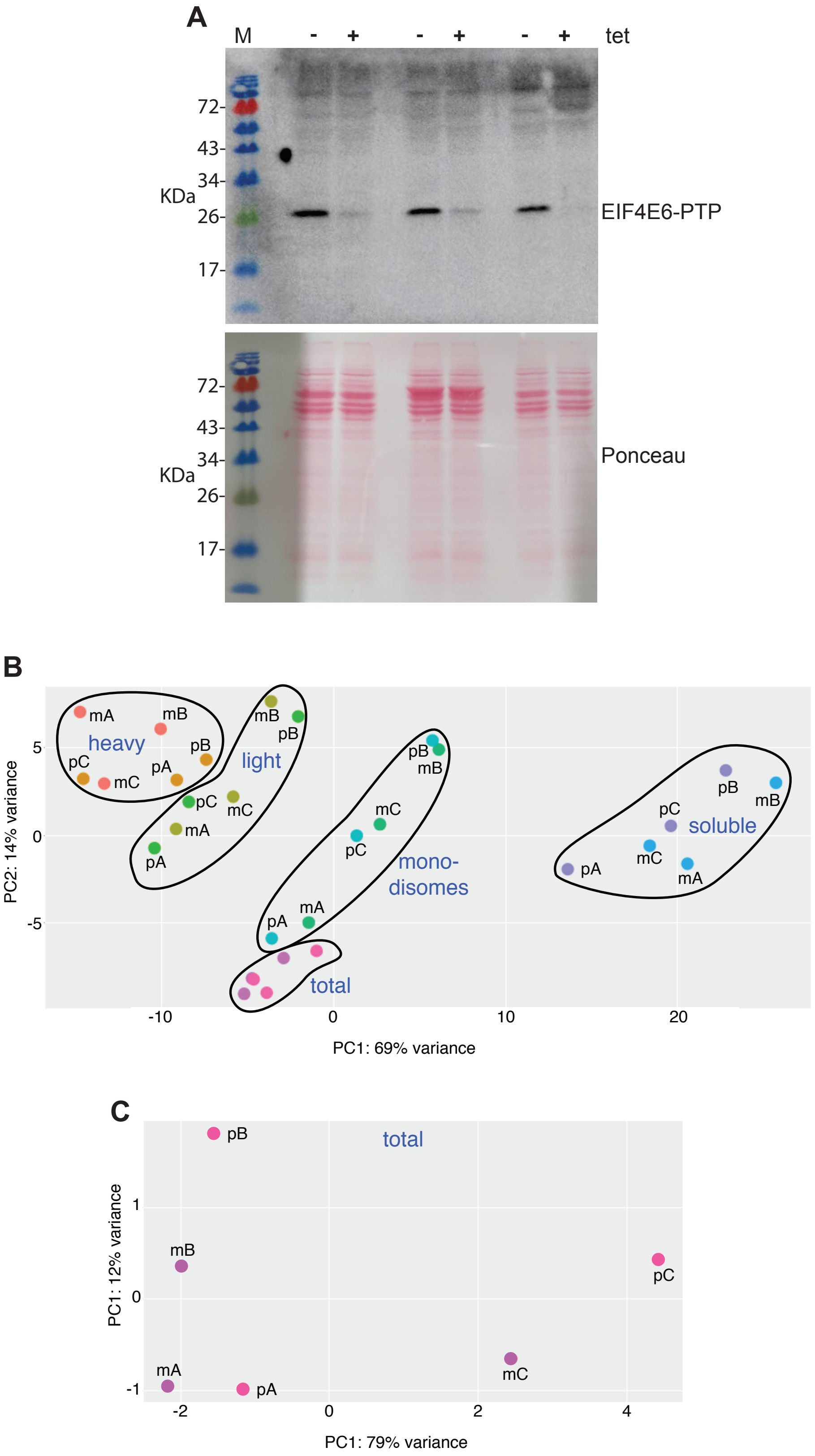

### Supplementary Figure S2

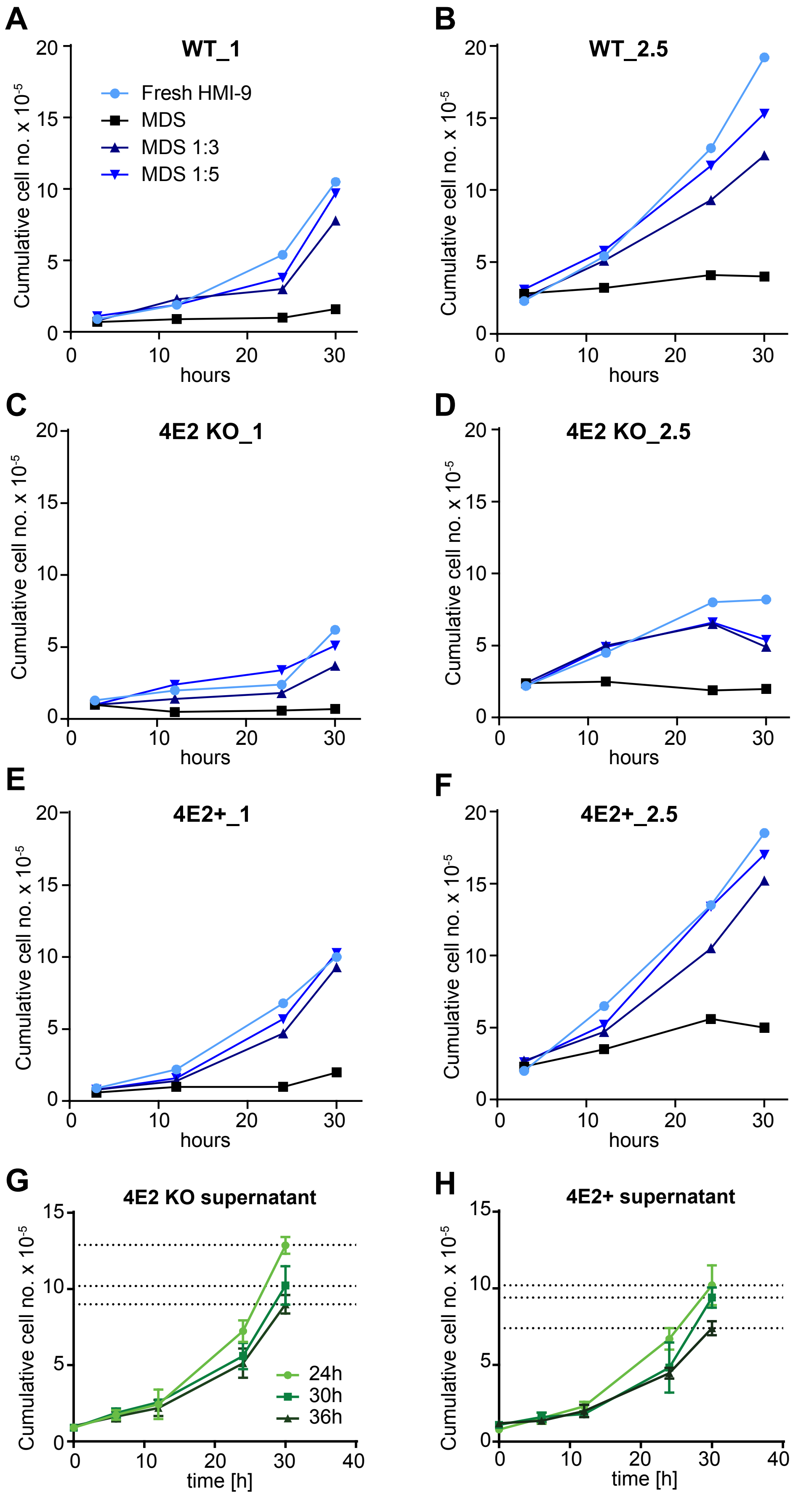

### Supplementary Figure S3

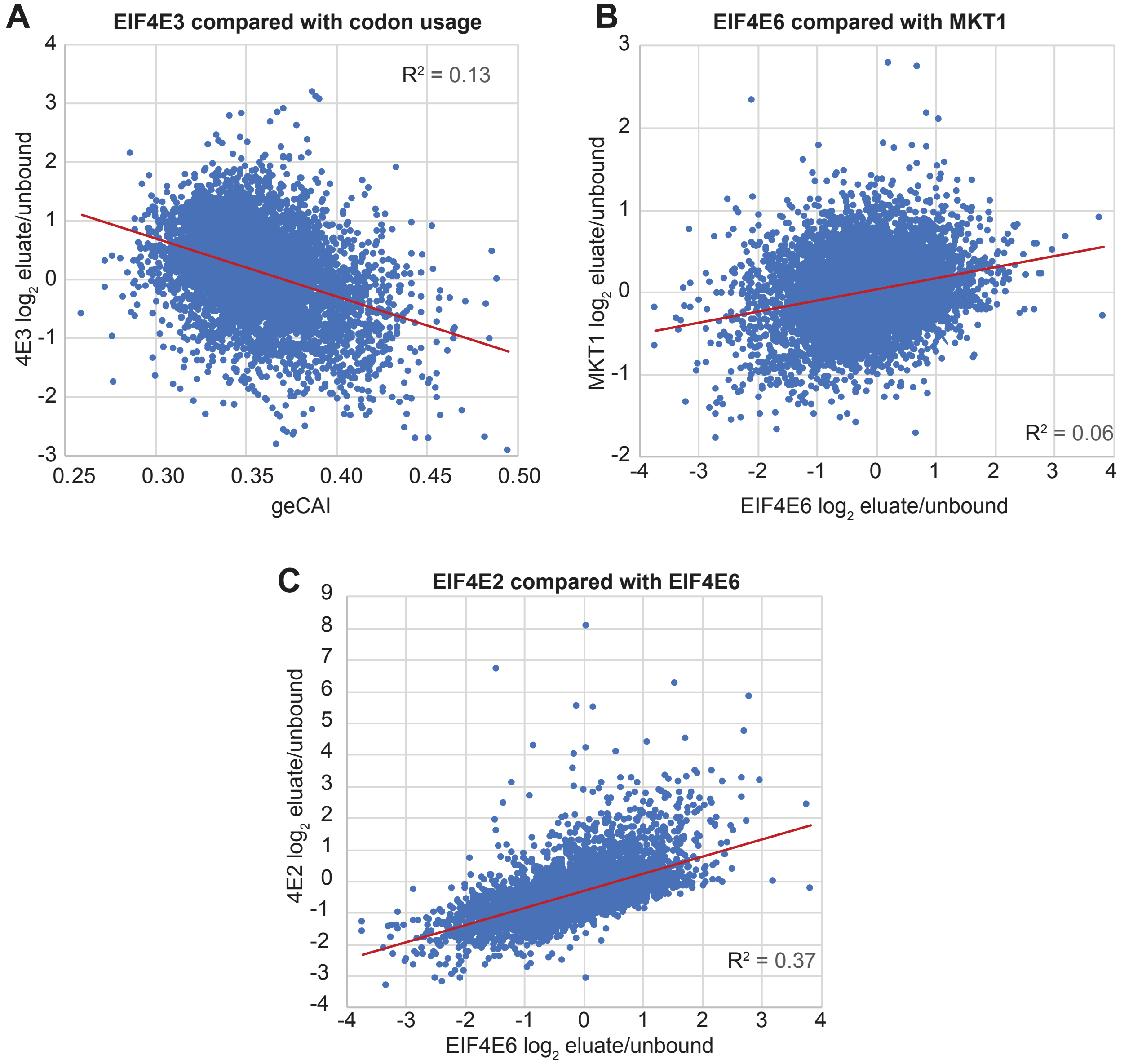

### Supplementary Figure S4

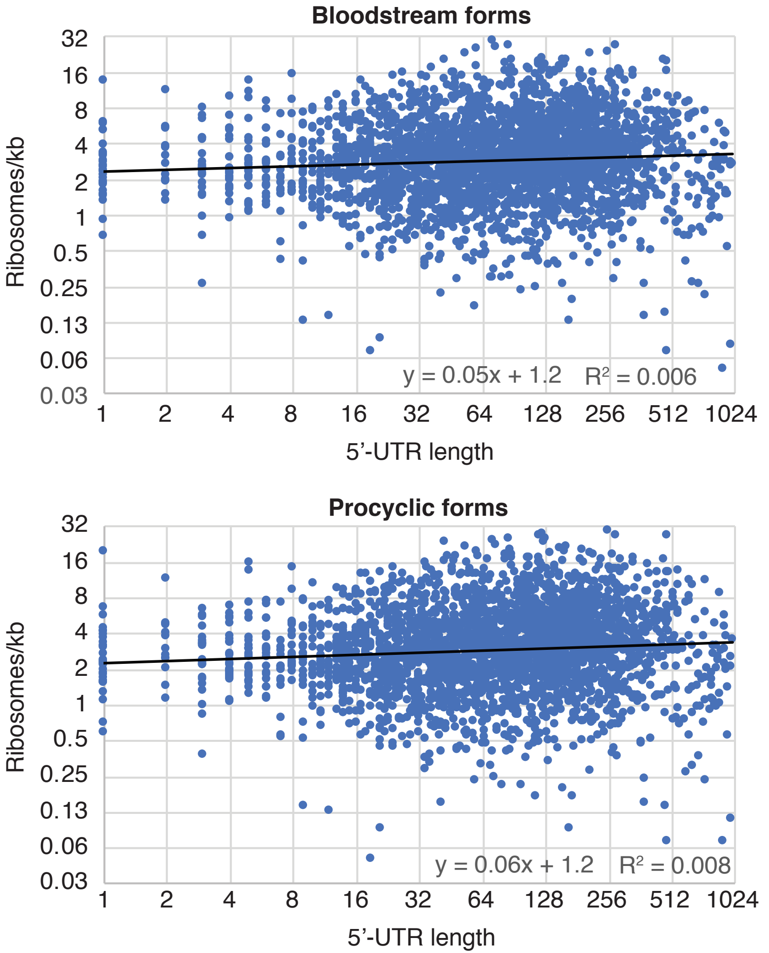
